# MixEHR-Guided: A guided multi-modal topic modeling approach for large-scale automatic phenotyping using the electronic health record

**DOI:** 10.1101/2021.12.17.473215

**Authors:** Yuri Ahuja, Yuesong Zou, Aman Verma, David Buckeridge, Yue Li

## Abstract

Electronic Health Records (EHRs) contain rich clinical data collected at the point of the care, and their increasing adoption offers exciting opportunities for clinical informatics, disease risk prediction, and personalized treatment recommendation. However, effective use of EHR data for research and clinical decision support is often hampered by a lack of reliable disease labels. To compile gold-standard labels, researchers often rely on clinical experts to develop rule-based phenotyping algorithms from billing codes and other surrogate features. This process is tedious and error-prone due to recall and observer biases in how codes and measures are selected, and some phenotypes are incompletely captured by a handful of surrogate features. To address this challenge, we present a novel automatic phenotyping model called MixEHR-Guided (MixEHR-G), a multimodal hierarchical Bayesian topic model that efficiently models the EHR generative process by identifying latent phenotype structure in the data. Unlike existing topic modeling algorithms wherein the inferred topics are not identifiable, MixEHR-G uses prior information from informative surrogate features to align topics with known phenotypes. We applied MixEHR-G to an openly-available EHR dataset of 38,597 intensive care patients (MIMIC-III) in Boston, USA and to administrative claims data for a population-based cohort (PopHR) of 1.3 million people in Quebec, Canada. Qualitatively, we demonstrate that MixEHR-G learns interpretable phenotypes and yields meaningful insights about phenotype similarities, comorbidities, and epidemiological associations. Quantitatively, MixEHR-G outperforms existing unsupervised phenotyping methods on a phenotype label annotation task, and it can accurately estimate relative phenotype prevalence functions without gold-standard phenotype information. Altogether, MixEHR-G is an important step towards building an interpretable and automated phenotyping system using EHR data.

## 1. Introduction

Electronic Health Records (EHRs) offer high volume comprehensive observational patient health data for clinical and translational research [1, 2]. The past 15 years have seen an explosion in EHR adoption, with the proportion of acute care hospitals and primary care practices in the United States using EHRs increasing from 9% and 17% respectively in 2008 to 96% and 86% by 2017 [3, 4]. Comprised of multiple types of data, such as lab tests, prescriptions, free-text clinical notes, and codified billing information including International Classification of Diseases (ICD), Current Procedural Terminology (CPT), and Diagnosis Related Group (DRG) codes, EHR data provide a comprehensive description of patients’ interactions with the healthcare system over time. Such rich, large-scale data have myriad potential applications including personalized disease risk estimation and treatment recommendation, pillars of precision medicine [5]. However, effective use of EHR data for such applications is often hampered by a lack of reliable disease phenotype labels.

Many studies use billing codes (i.e. ICD codes) as surrogates for phenotype labels, a practice that works well for some phenotypes but notoriously poorly for others, including rheumatoid arthritis and chronic kidney disease [6–9]. Other studies have physicians or other health experts manually review patient records or devise rule-based phenotyping algorithms based on surrogate features such as ICD codes [10–17]. An example of this approach is the PheKB community phenotyping knowledge base, which has shown considerable success in generating accurate, portable rules [17]. While generally reliable, both chart review and rule production are tedious, time-intensive processes applicable to labeling a handful of phenotypes for discovery research but not for phenotyping at scale, as might be needed for a multi-phenotype hypothesis-generating study such as a Phenome-Wide Association Study (PheWAS). Moreover, rule-based methods are subject to recall and observer biases in how features are selected.

To address this challenge, researchers have proposed a variety of automated computational phenotyping algorithms that infer phenotypes with little to no expert input or annotation. Harutyunyan et al. (2019) [18] used 17 handcrafted clinical variables to classify patients to 25 Clinical Classifications Software (CCS) codes using logistic regression or recurrent neural networks, with each CCS defining a group of related ICD-9 codes as a disease category. Some notable unsupervised learning approaches include work in non-negative matrix/tensor factorization [19–21], mixture models [22–30], and neural network-based embedding learning [18, 31, 32]. One strategy that has shown considerable success is to use a handful of quickly identifiable surrogate features, including ICD codes and natural language processing (NLP)-derived mentions of a target phenotype in clinical notes, to generate “silver-standard” phenotype labels - or “anchors” - that can be subsequently used to guide phenotype estimation in a “weakly supervised” manner [25, 26, 29, 30, 33]. An ongoing challenge remains how best to leverage the thousands of additional features in the EHR to improve upon these silver-standard labels. Several methods employ regression with regularization or dropout to de-noise silver-standard labels using other features [25, 33], but this strategy tends to be sensitive to high feature dimensions and sparsity typical for EHR data.

Topic modeling algorithms are well-conditioned to this setting. Designed to model highly-sparse text data with vocabularies often consisting of tens of thousands of words, topic models have shown considerable aptitude at discovering latent structure (i.e. topics) in such datasets [30, 34–36]. Indeed, several studies have demonstrated success applying derivatives of the widely used topic model Latent Dirichlet Allocation (LDA) to EHR data, considering patients as documents, disease phenotypes as topics, and EHR features as words [30, 34]. One notable model, MixEHR, builds upon LDA to 1) simultaneously model an arbitrary number of data modalities (e.g., ICD codes, lab results) with modality-specific distributions, and 2) treat observations as not missing at random (NMAR), reflecting the fact that clinicians typically collect data such as lab tests with a diagnosis in mind [34]. However, MixEHR is fully unsupervised and thus prone to learning topics that lack clinical meaning. On the other hand, the recent sureLDA method constrains topics to align with well-defined phenotypes using a weakly supervised surrogate-based approach, thereby learning meaningful, interpretable topics [30]. However, sureLDA treats all features as draws from topic-specific multinomial distributions, rendering it unable to properly model other data modalities such as lab tests and vital signs. Moreover, its Gibbs sampling inference procedure does not scale well to datasets with more than thousands of features or a few hundred thousand patients.

In this study we present MixEHR-Guided (MixEHR-G), which is built upon two previously established methods: sureLDA [30] and MixEHR [34]. Our key contribution is the elegant use of guided topic priors originally introduced in sureLDA to efficiently infer multi-modal topic distributions over heterogeneous EHR data types using a variational inference algorithm, which was introduced in MixEHR. This powerful combination allows us to infer from large-scale EHR datasets over 1500 multi-modal phenotype topics each corresponding to a unique well-defined disease phenotype. We apply MixEHR-G to the Population Health Record (PopHR) dataset [37], which contains longitudinal administrative claims data for a random sample of 1.3 million residents of Quebec, Canada (**Methods**), and to the Medical Information Mart for Intensive Care (MIMIC-III) dataset [38], a publicly available dataset of Intensive Care Unit (ICU) observations at the Beth Israel Deaconess Medical Center in Boston, Massachusetts. In both applications, MixEHR-G is able to effectively infer meaningful phenotype distributions over high-dimensional multimodal categorical EHR data including ICD codes, prescriptions, and discretized lab results. We use MixEHR-G’s patient-topic mixture distributions to predict patient phenotypes and to analyze disease prevalence as a function of age and sex.

## 2. Methods

### 2.1. MixEHR-G model description

Building upon MixEHR, MixEHR-G uses a probabilistic joint topic modeling approach that projects a patient’s high-dimensional and heterogeneous clinical record onto a low-dimensional latent topic mixture membership over disease phenotypes. In contrast to MixEHR, MixEHR-G incorporates noisy phenotype labels obtainable *a priori* as topic priors for each phenotype to guide posterior topic inference. In this way, MixEHR-G harnesses the unsupervised topic modeling prowess of MixEHR to learn interpretable and well-defined phenotypes in a manner similar to the anchor-and-learn framework [39] or sureLDA [30]. In this study, we use Phenotype Codes (PheCodes), expert-defined groupings of ICD-9 codes developed for Phenome-Wide Association Studies (PheWAS), as topic priors. PheCodes have been shown to better align with clinically meaningful disease phenotypes and better replicate known genetic associations in genome-wide association studies (GWAS) than individual ICD-9 codes or CCS groupings of ICD codes [40]. That said, MixEHR-G will work with any phenotype prior, including the ICD taxonomy or CCS codes; PheCodes are not the only phenotypic anchor schema compatible with our method. Crucially, the only change to MixEHR’s computational framework that MixEHR-G introduces is the PheCode-driven initialization and re-optimization of the Dirichlet hyperparameter *α*, so no functionality featured by MixEHR is lost.

The MixEHR-G algorithm consists of 3 key steps: (1) assemble a comprehensive and heterogeneous EHR dataset (i.e. ICD codes, prescriptions, lab data, etc.); (2) use the Multimodal Automated Phenotyping (MAP) method to obtain initial probabilities for each reference phenotype based on key surrogate features within these data; and (3) train the MixEHR model [34] on the entire EHR dataset weighting the *K* asymmetrical patient-topic Dirichlet hyperparameters by the probabilities obtained in (2). We assume that there are in total K reference phenotypes corresponding to K true binary phenotype states that are unobserved *a priori*. Given a patient’s EHR data, we predict the K phenotypes by inferring the posterior distributions of K guided topics, each corresponding to exactly one reference phenotype due to the use of constrained topic priors. **Fig.1** depicts this procedure, and we now describe each of the 3 steps in detail.

**Figure 1:**
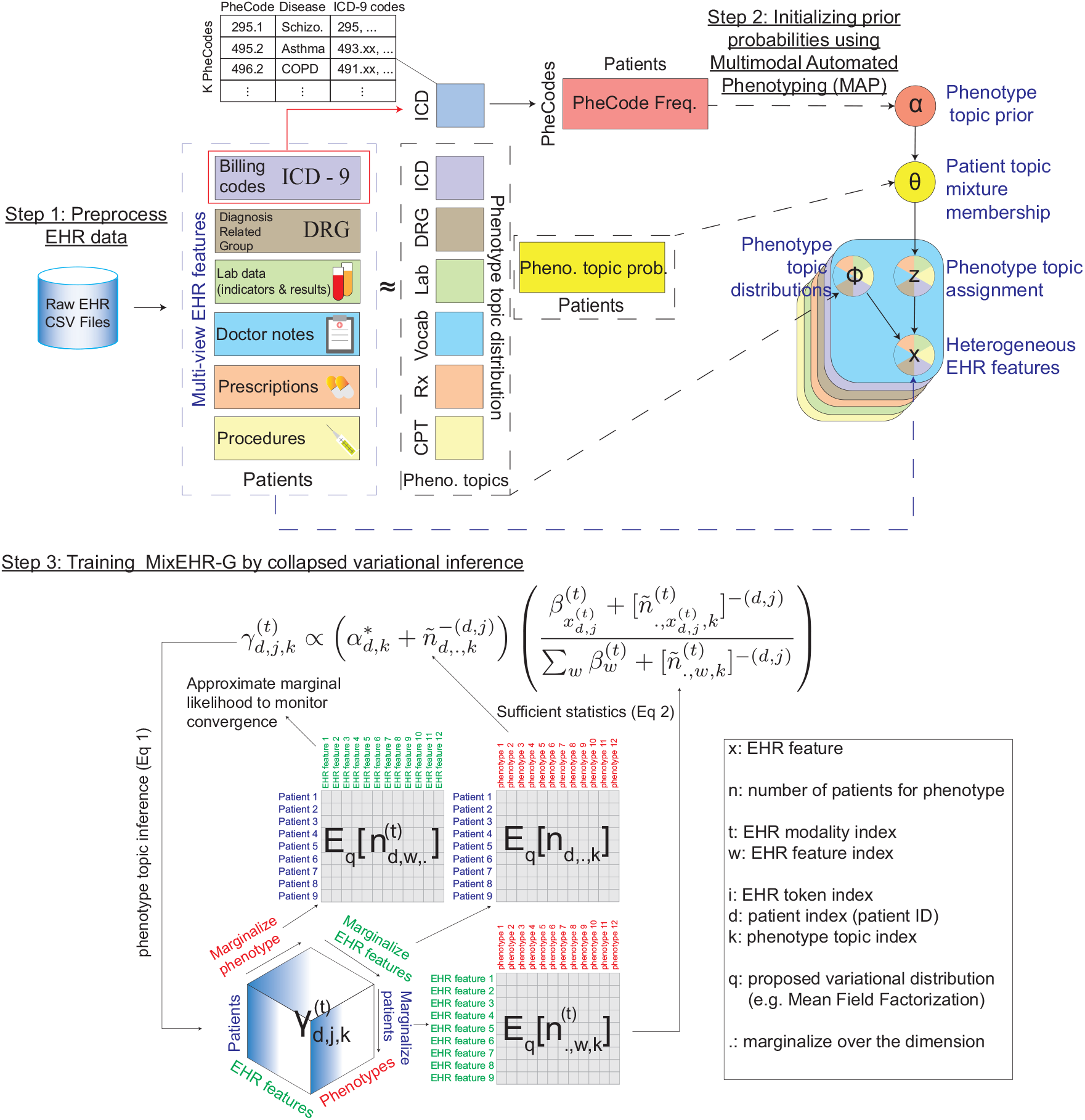
Schematic of the MixEHR-G algorithm. We factorize the heterogeneous EHR data matrices into phenotype topic distributions (i.e. phenotype topic distribution matrices) and 1 common patient topic mixture (i.e. patient phenotype probability matrix). To guide patient topic inference, we first infer the PheCode-guided topic prior ***α**_d_* for each patient *d*. This is done by training Multimodal Automated Phenotyping (MAP), a two-component Lognormal and Poisson Mixture models on the normalized frequency of the ICD codes that define the PheCode, and taking a weighted average of the estimated posterior probabilities for the higher components of the two models. We then incorporate this weighted posterior as the Dirichlet hyperparameters ***α**_d_* for the patient topic mixture θ_*d*_. Finally, we infer the posterior distributions of the phenotype topic assignment for each EHR token *j* of patient *d* (*z_d,j_*) and phenotype topic distributions (ϕ) using a collapsed mean-field variational inference algorithm. Each of the 3 steps are described in detail in **Methods** Section 2.1. Step 3 highlights the key algorithmic step to train MixEHR-G, which alternates between topic inference (Eq 1) and sufficient statistics (Eq 2) computation.

#### Step 1: preprocess EHR data

Following a topic modeling approach [41], we treat each patient record as a document and the various EHR features throughout that patient’s record - including all CPT codes, DRG codes, ICD codes, RxNorm codes, clinical notes, and lab data - as tokens. Because we do not consider the sequential order of observed codes, we essentially model the patient record as a bag of words (or rather a bag of EHR features). EHR features are often filtered by feature selection methods such as the Surrogate-Assisted Feature Extraction (SAFE) method [28], which may filter out useful information. Instead MixEHR-G has the capacity to assign probability weights to each EHR feature under each phenotype topic, making it robust to a large number of uninformative codes. We included all available features without any filtering. **Fig.1** depicts how multimodal count data are combined into multiple discrete-count matrices - one per data type - for input into the MixEHR-G algorithm.

#### Step 2: Initializing prior probabilities using MAP

We obtain prior probabilities for the *K* reference phenotypes, which we denote as ***π*** = (*π*_1_,…, *π_K_*)’, using a modified MAP algorithm [26]. Standard MAP estimates *π_k_* by fitting Poisson and Lognormal mixture models to the counts of (1) phenotype *k*’s core ICD code and (2) NLP-curated mentions of phenotype k in clinical notes, normalizing using a healthcare utilization feature *H*, and taking a weighted average of these various mixture models. Instead of ICD and NLP features, we use Phenotype Codes (PheCodes), which are coarser than ICD codes and have been found to better align with diseases described in clinical practice [40]. We mapped ICD codes to PheCodes using the established PheWAS mapping [42]. We then compiled PheCode counts at the level of (1) integer codes (i.e. 496.21 →496 (COPD)), which we refer to as parent phenotypes, and (2) 1-decimal codes (i.e. 496.21 →496.2 (chronic bronchitis)), which we refer to as subphenotypes. We then ran MAP (including both Poisson and Lognormal mixture models) on these phenotype and subphenotype PheCode counts to estimate prior probabilities for each corresponding (sub)phenotype. For most diseases, patients with a corresponding PheCode count of 0 (i.e. filter-negative) rarely have the disease, so we set the corresponding topic prior *π_k_* = 0 for these patients. Note that we do not include additional ‘empty’ topics to model information not contained in the PheCodes as we assume that including a topic for every PheCode sufficiently covers all phenotype information in each dataset.

#### Step 3: training MixEHR-G

MixEHR models EHR data using a generative latent topic model inspired by Latent Dirichlet Allocation [34, 41] (**Fig.1)**. We assume that each phenotype topic *k* ∈ {1,…, *K* } under EHR data type *t* ∈ {1,…, *T*} is characterized by a latent probability vector over *W*^(*t*)^ EHR features, 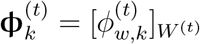, which follows a Dirichlet distribution with unknown hyperparameter *β_wt_*. For each patient *d* ∈ {1,…, *D*}, we assume that the patient’s phenotype mixture membership ***θ**_d_* is generated from a K-dimensional Dirichlet distribution with unknown asymmetric hyperparameters ***α*** = [*α_k_*]_*K*_. Thus, to generate EHR observation *j* of datatype *t* for patient *d*, we first draw a latent topic 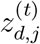 from a multinomial distribution with probability vector **θ**_*d*_. Given 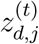, we then draw an EHR feature 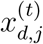 from a multinomial distribution with probability vector 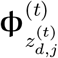.

MixEHR-G extends MixEHR by setting the Dirichlet topic hyperparameters for patient *d*, ***α**_d_*, to a scalar multiplied by the prior probability vector over the *K* phenotypes: ***α**_d_* = *α**π**_d_*, where ***π**_d_* is obtained from Step 2. Consequently, patient *d*’s expected mixture membership for phenotype *k, α_d,k_* is scaled by patient *d*’s prior probability of phenotype *k*, reflecting the intuition that a higher prior likelihood of phenotype presence should beget a proportionally higher probability of feature attribution to that phenotype. Therefore, patient *d* will have zero mixture membership for phenotype *k* if the topic prior is equal to zero (*π_d,k_* = 0).

We train MixEHR-G using the joint collapsed variational Bayesian inference protocol detailed in Li et al. (2020) [34]. The key variational topic inference step is:

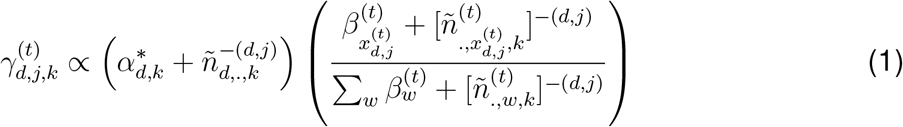

where 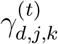 denotes the variational probabilities of the *k^th^* topic assigned to feature *j* of patient *d*: 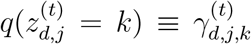 and the notation *n*^−(*d,j*)^ indicates the exclusion of the contribution from patient *d* and feature *j*. The sufficient statistics are simply the summation of the inferred topic probabilities over all patients and all features except for patient *d* and feature *j*:

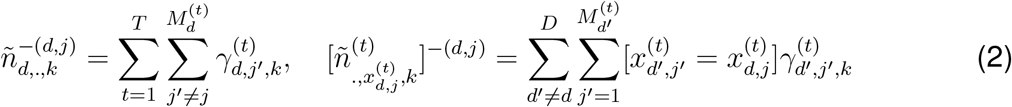

Unlike MixEHR, instead of iteratively optimizing the asymmetrical hyperparameters *α*, we now just optimize the scalar *α*. During the M-step of the EM learning algorithm, we update *α* by maximizing the marginal likelihood under the variational expectation via an empirical Bayes fixed-point update:

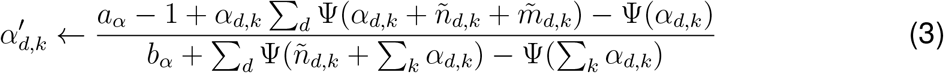

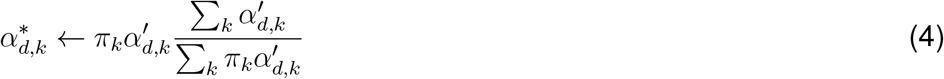

The remainder of the algorithm is identical to that detailed in Li et al. (2020) [34] and thus omitted here. All functionality featured by MixEHR-G, including imputation of missing lab results, is preserved. When MixEHR-G converges, we obtain the count of total EHR features assigned to phenotype topic *k* for subject *d* as the patient-level phenotype mixture score: 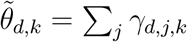. We use 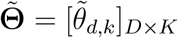 for *D* patients over *K* topics for downstream analyses.

### 2.2. Implementation of existing methods

We implemented sureLDA [30], MAP [26], and PheNorm [25] using their respective R packages: sureLDA:https://cran.r-project.org/web/packages/sureLDA/index.html; MAP:https://search.r-project.org/CRAN/refmans/MAP/html/MAP.html; Phe-Norm:https://cran.r-project.org/web/packages/PheNorm/index.html. We implemented MixEHR [34] using the source code from the github page:https://github.com/li-lab-mcgill/mixehr.

## 3. EHR datasets

### 3.1. EHR data overview

We applied MixEHR-G with phenotype topics defined by the set of *K* unique PheCodes (including both integer-level and 1-decimal codes in the PheCode hierarchy) to the PopHR and MIMIC-III datasets [37, 38]. Briefly, PopHR contains both chronic and acute disease information from outpatient visits as recorded in ICD codes, administrative treatment (ACT) codes, and RxNorm medication codes. MIMIC-III contains highly acute disease information from ICU admissions, often with comorbid or causative chronic diseases (e.g., flash pulmonary edema secondary to heart failure in a patient with chronic hypertension), as recorded in clinical notes, ICD-9 billing codes, prescriptions, DRG billing codes, CPT procedural codes, lab tests, and lab results. We identified *K =* 1387 PheCodes in PopHR (512 3-integer codes and 875 3-integer+1-decimal codes) and K = 1515 in MIMIC-III (499 3-integer codes and 1016 3-integer+1-decimal codes), reflecting the different clinical settings of the two datasets. Reassuringly, we found that patients tend to have multiple PheCode anchors in both datasets (a median of 14 integer codes and 16 1-decimal codes per patient in PopHR, and a median of 6 integer codes and 7 1-decimal codes per patient in MIMIC-III) **Supplementary Fig.** S1. This suggests that the PheCodes are sensitive but not overly specific, providing strong guidance for MixEHR-G’s topics while allowing for learning and divergence from the anchor. We used these two datasets to demonstrate the generalizability of MixEHR-G and gain contrasting insights between the datasets regarding the characterization of a common set of diseases. Besides the distinct population coverage of the two datasets, they also involve different feature modalities and levels of disease acuity (**Methods**).

Using MIMIC-III data as a demonstration (**Fig.1)**, we learned seven sets of topic distributions in the form of the basis matrices corresponding to each of the data modalities detailed above. Notably, lab data contribute two modalities corresponding to the binary indicators (i.e, whether the lab test was ordered) and lab test results coded as categorical variables with discrete values of normal and abnormal (more details below). We link these seven basis matrices using a common patient-topic mixture matrix, which is proportional to the EHR data multinomial likelihood and the prior inferred from the ICD-9 signatures under each PheCode.

### 3.2. Applications to the MIMIC-III dataset

MIMIC-III is a large, single-center database comprising multimodal EHR data associated with 53,423 distinct hospital admissions for 38,597 adult patients and 7,870 neonates admitted to critical care units at the Beth Israel Deaconess Medical Center in Boston, MA between 2001 and 2012 [38]. The dataset was downloaded from the PhysioNet database (mimic.physionet.org) and used in accordance with the PhysioNet user agreement.

We used the same preprocessing pipeline as in [34]. Specifically, clinical notes from `NOTEEVENTS.csv` were pre-processed and converted to bag-of-words format using the `R` library `tm`. We filtered out common stop words, punctuations, numbers, and whitespace, and we converted all remaining words to lowercase.

For lab data from `LABEVENTS.csv`, we utilized the FLAG column to ascertain whether a test result was normal or abnormal. For a given patient, we recorded the frequencies of both normal and abnormal lab results for each test.

For prescription data from `PRESCRIPTIONS.csv`, we concatenated the `DRUG_NAME_GENERIC`, `GSN`, and `NDC` columns to create a compound ID for each prescription. We combined entries with the same compound ID to eliminate feature redundancy by drug formulation. Likewise, for diagnosis-related group (DRG) codes from `DRGCODES.csv`, we created a compound ID as the concatenation of `DRG_TYPE`, `DRG_CODE`, `DESCRIPTION`, `DRG_SEVERITY`, and `DRG_MORTALITY`. We kept the original IDs for ICD-9 diagnosis codes (`ICD-9-CM`) from `DIAGNOSES_ICD.csv` and procedure codes (`ICD-9-CPT`) from `PROCEDURES_ICD.csv`. For each data type, we removed nonspecific features that were frequently observed among patients based on the inflection point at the curve of the increasing feature frequencies (i.e., all the features below the inflection point were retained as shown in Supplementary Figure 26 in [34]).This resulted in 33,388 unique words from clinical notes, 6,215 ICD-9 codes, 1,770 CPT codes, 564 lab tests, 8,409 prescriptions, and 3,086 DRG codes, altogether totaling 53,432 unique features.

The MIMIC-III data are not longitudinal. Most patients in the MIMIC-III dataset only have one ICU admission, wherein each ICD-9 code is observed at most once upon discharge. As a result, each PheCode is observed at most once for each patient. In lieu of running MAP on PheCode counts to obtain prior phenotype probabilities as detailed in *Step 2: Initializing prior probabilities using MAP*, we set the hyperparameters *α_d,K_* for phenotype *k* to one or zero depending on whether the associated PheCode was observed for patient *d* or not, respectively.

### 3.3. Applications to the PopHR dataset

PopHR is a multimodal database that integrates massive amounts of longitudinal heterogeneous data from multiple distributed sources (e.g. inpatient and outpatient physician claims, hospital discharge abstracts, outpatient drug claims) for 1.3 million patients in the Quebec province of Canada between 1998 and 2014 [37]. We used three data modalities from this dataset: ICD-9 diagnostic codes, administrative procedure codes (ACT), and prescriptions all from outpatient physician encounters. For prescriptions, we used drug identification numbers (DINs) as unique IDs; for the other two we used ICD-9 and ACT codes respectively. We removed all features with patient frequencies above 25%to mitigate the influence of common, imprecise features. This left 5,935 unique ICD-9 codes, 4,037 ACT codes, and 1,311 DIN codes for a total of 11,283 unique count features. Some DIN codes indicate different dosages or formulations of the same active drug ingredient. We found this to be useful information for modeling phenotypes as the same drug is often prescribed differently for different conditions. However, for qualitative topic analyses such as the heatmap in **Fig.2a**, we aggregated topic probabilities for DIN codes by active drug ingredient to simplify interpretation.

**Figure 2:**
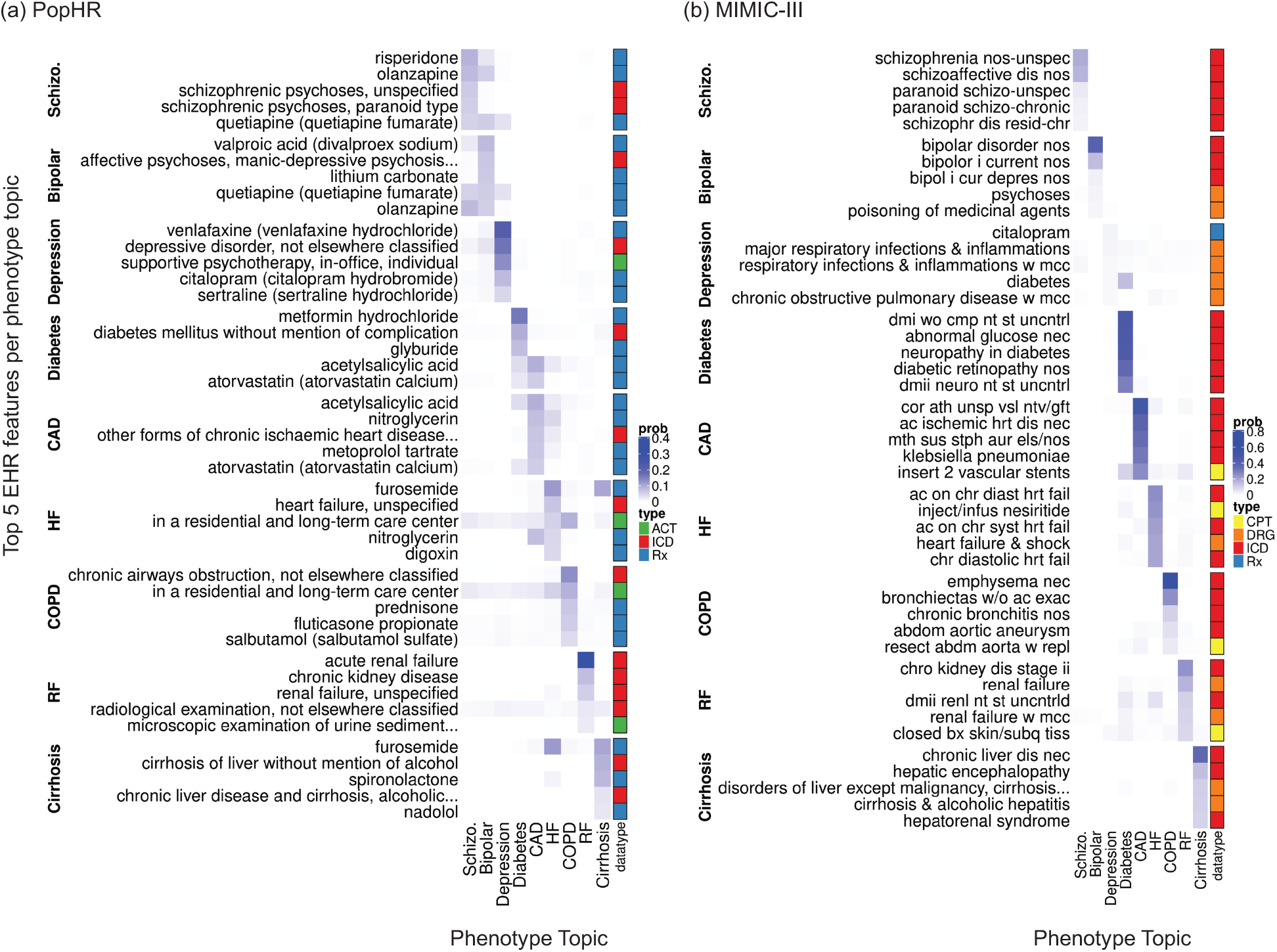
Top 5 features for each of 9 diverse disease phenotypes. (**a**) Phenotype topic distributions inferred from the PopHR dataset. Using MixEHR-G, we inferred phenotype topic distributions (ϕ) for all PheCode-guided parent phenotypes and subphenotypes and displayed 9 of them. We include distributions for all phenotype topics in the Supplementary Materials. Phenotype topic distributions are inferred over EHR features of 3 data types (ICD, ACT, Rx). (**b**) Phenotype topic distributions inferred from the MIMIC-III data. Phenotype distributions were inferred over EHR features of 6 data types (CPT, DRG, ICD, Rx, clinical notes, and lab data). Heatmap color intensities reflect the probabilities of features under a given phenotype. The color bar on the right side of each heatmap indicates the data types as shown in the legend. Only data types corresponding to the top EHR features under each phenotype are visible in the heatmaps. Some row names were truncated due to length. In panel (a) the truncated names are ‘affective psychoses, manic-depressive psychosis, other and unspecified’, ‘other forms of chronic ischaemic heart disease, unspecified’, ‘microscopic examination of urine sediment and interpretation’, ‘chronic liver disease and cirrhosis, alcoholic cirrhosis of liver’; in panel (b) the truncated name is ‘disorders of liver except malignancy, cirrhosis, alcoholic hepatitis with complications, comorbidities’. The full names for the abbreviated column names are as follows: Schizo.: Schizophrenia; Bipolar: Bipolar Disorder; CAD: Coronary Artery Disease; HF: Heart Failure; COPD: Chronic Obstructive Pulmonary Disease; and RF: Renal Failure.

Unlike MIMIC-III, PopHR is a longitudinal dataset in which a subject can encounter multiple occurrences of any given ICD-9 code (and thus PheCode) over time. Thus, we computed prior phenotype probabilities using the MAP procedure described in *Step 2: Initializing prior probabilities using MAP*.

## 4. Experimental design

### 4.1. Phenotyping accuracy evaluation

For the PopHR data [37], we applied expert-defined rule-based phenotyping algorithms for 12 chronic diseases. We selected these 12 diseases - schizophrenia, ischemic heart diseases (IHD), hypertension, HIV, epilepsy, diabetes, COPD, congestive heart failure (CHF), autism, asthma, acute myocardial infarction (AMI), and ADHD - as they were the only phenotypes with clear PheCode alignments for which we had gold-standard rule-based algorithms. For 9 of these phenotypes - acute myocardial infarction (AMI), asthma, congestive heart failure (CHF), COPD, diabetes, hypertension, ischemic heart disease (IHD), epilepsy, and schizophrenia - rules were defined by the Chronic Disease Surveillance Division of the Public Health Agency of Canada [43]. For the remaining 3 phenotypes, for ADHD we used the definition described in Vasiliadis et al. (2017) [44], for autism we followed the 2017 Autism Spectrum Disorder Surveillance in Quebec report [45], and for HIV we applied the definition described in Durand et al. (2011) [46].

All rules are derived entirely from ICD codes, sometimes relying on consecutive observations of the same ICD code(s) for a patient within a small (i.e. 1 month) time interval. Precise definitions for each rule-based phenotype can be found in the **Supplementary Materials**. Since we aggregated ICD code frequencies over time for a given patient, our model does not use the same temporal information as the rules, making them a good reference for MixEHR-G. Therefore, we used the phenotype labels derived from the rules as gold-standards to evaluate our unsupervised model performance. We do not apply these rule-based phenotyping algorithms to the MIMIC-III dataset as they were intended to identify chronic diseases using longitudinal data, whereas MIMIC-III only contains data for hyperacute ICU visits.

For both the MIMIC-III and PopHR datasets, we ran MixEHR-G on parent phenotypes and subphenotypes separately. We then derived the MixEHR-G phenotype score 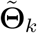 for a coarse phenotype such as COPD (496) using the corresponding topic from the parent phenotype model, whereas for subphenotypes such as emphysema (496.2) we used the subphenotype model.

### 4.2. Identification of similar and comorbid phenotypes

We defined the *similarity* between two phenotypes *k* and *k*’ as the Spearman correlations between their respective MixEHR-G topic distributions ϕ_*k*_ and ϕ_*k*_’ over the *V* unique EHR features across data modalities. We defined the *comorbidity* between phenotypes *k* and *k*’ as the Spearman correlations between their respective patient mixture proportions **θ**_*k*_ = [*θ_d,k_*]_*D*_ and **θ**_*k*_’ = [*θ_d,k_*’]_*D*_. These choices reflect the intuition that two phenotype topics with similar distributions over EHR features are qualitatively similar, whereas two topics with similar mixture probabilities over patients are comorbid. Thus, for each phenotype of interest, we use the trained **ϕ**_*K×V*_ matrix to identify similar *parent* phenotypes, and we use the trained **θ**_*D×K*_ matrix to identify comorbid *parent* phenotypes.

### 4.3. Estimation of relative population phenotype prevalence stratified by age and sex

For a given phenotype *k*, we set the overall phenotype prevalence *ρ_k_* to the estimate from the Global Health Data Exchange, a comprehensive catalog of health-related data compiled by the Institute for Health Metrics and Evaluation (IHME) at the University of Washington [47]. We sought to estimate the relative prevalence stratified by age and sex using the PopHR dataset, which is representative of the general population (**Fig.7)**. First, we set a threshold 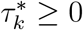 to the patient-topic mixtures 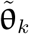 to derive predicted phenotype outcomes 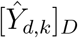 with mean value equal to the known population prevalence *ρ_k_*. In other terms, we denote the empirical quantile function of 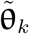 as 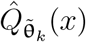,

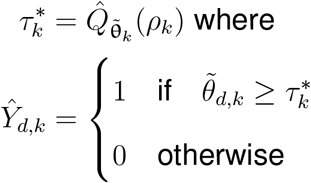

**Figure 3:**
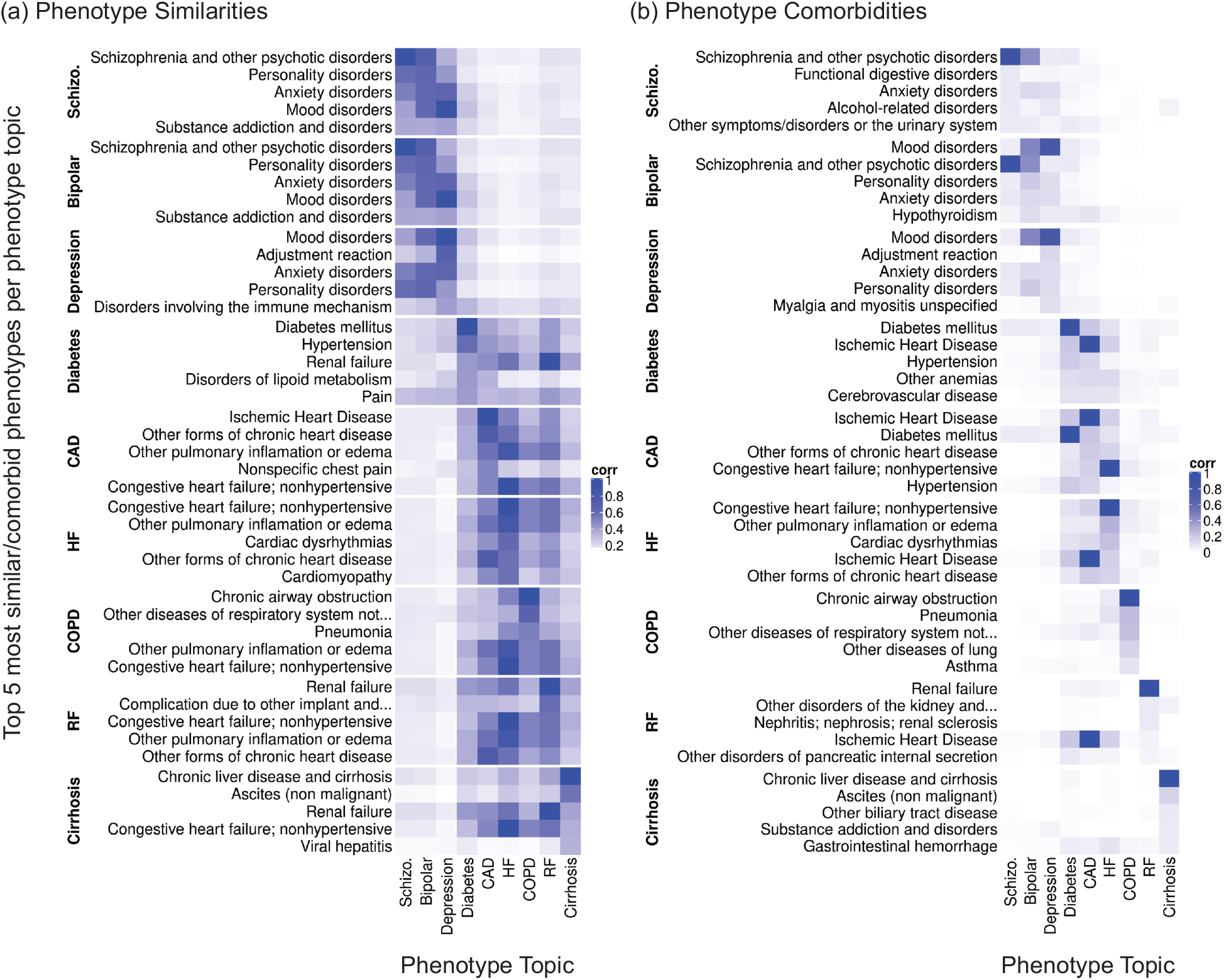
Predicted similarities and comorbidities of 9 diverse phenotypes. (**a**) 5 most similar parent phenotype topics for each of 9 diverse disease phenotypes, as predicted by *post-hoc* Spearman correlations among MixEHR-G’s topic distributions **Φ** over all multimodal EHR features. (**b**) 5 most comorbid phenotype topics per target disease as predicted by *post-hoc* Spearman correlations among MixEHR-G’s patient-topic mixtures 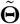. MixEHR-G was trained on the PopHR dataset. Heatmap color intensities are proportional to the Spearman correlation values. The color bar on the right side of each heatmap indicates the data types as shown in the legend. Only data types corresponding to the top EHR features under each phenotype topic are visible in the heatmaps.

**Figure 4:**
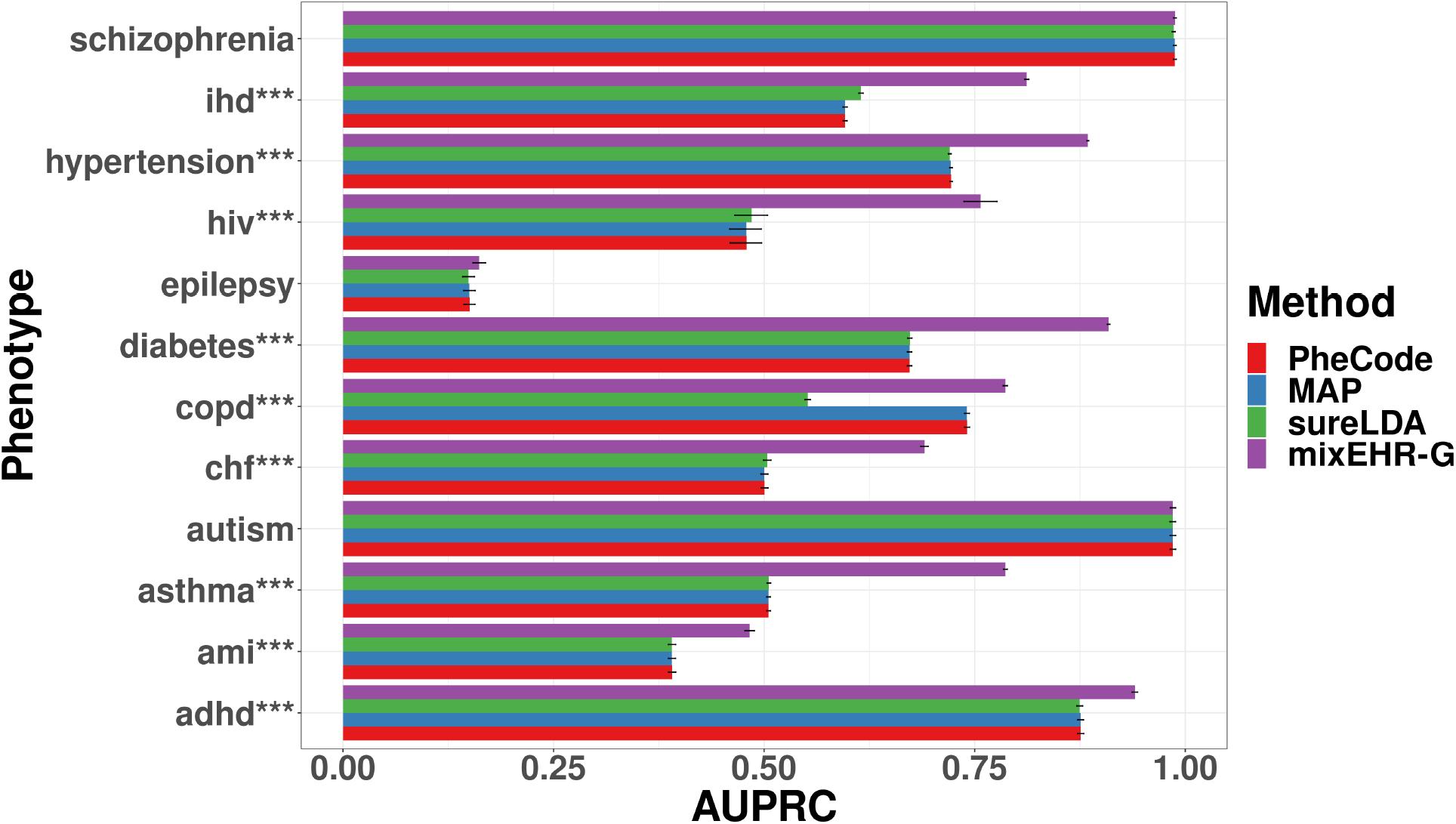
Predictive performance of 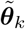 for phenotypes with expert-derived rule-based labels. We applied MixEHR-G and two recently published unsupervised phenotyping methods, MAP and sureLDA, to the entire PopHR data without supervision or feature filtering. We also computed the frequency of the raw PheCode counts for each target disease. The barplots display the areas under the precision recall curve (AUPRCs) within the PopHR dataset taking rule-based phenotype labels as gold-standards. 95%confidence intervals and standard errors were estimated by bootstrapping with 1000 subsamples. P-values were estimated using a two-sample t-test with bootstrapped standard error estimates, and the largest (i.e. least significant) p-value between MixEHR-Gand comparators for each target disease is displayed here. (*): p < 0.05; (**): p < 0.01; (***): p < 0.001.

**Figure 5:**
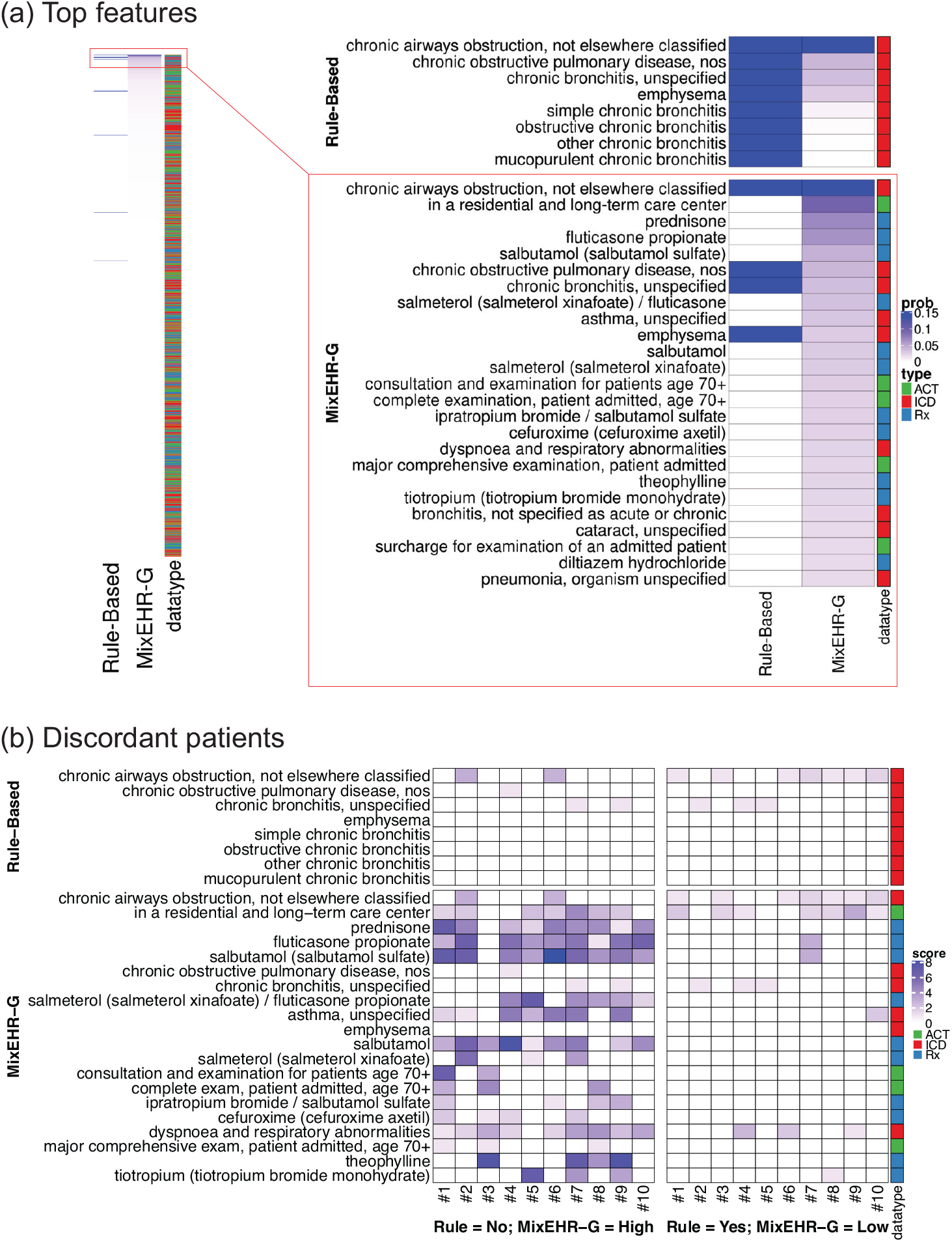
In-depth analysis of MixEHR-G’s phenotyping results for chronic obstructive pulmonary disease (COPD) using the PopHR dataset. (**a**) All 11,283 PopHR features (left) and the top 25 EHR features (right) with corresponding data types (color coded) per the COPD topic distribution ϕ_COPD_. We compared these features with the features included in the expert-curated rule-based COPD algorithm obtained from the Chronic Disease Surveillance Division of the Public Health Agency of Canada [43] described in the **Methods**. Features included in the rule-based algorithm are shown in the heatmap on top. (**b**) Profiles of the 20 most discordant patients. The heatmap on the left displays the 10 patients for whom MixEHR-G predicts the highest mixture scores for the COPD topic 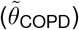 but the rule-based algorithm predicts them as negative. The heatmap on the right displays the 10 patients for whom MixEHR-G predicts the lowest COPD topic mixture scores but the rule-based algorithm predicts them as positive. The color intensity indicates the frequency count of each EHR feature for the selected patients.

**Figure 6:**
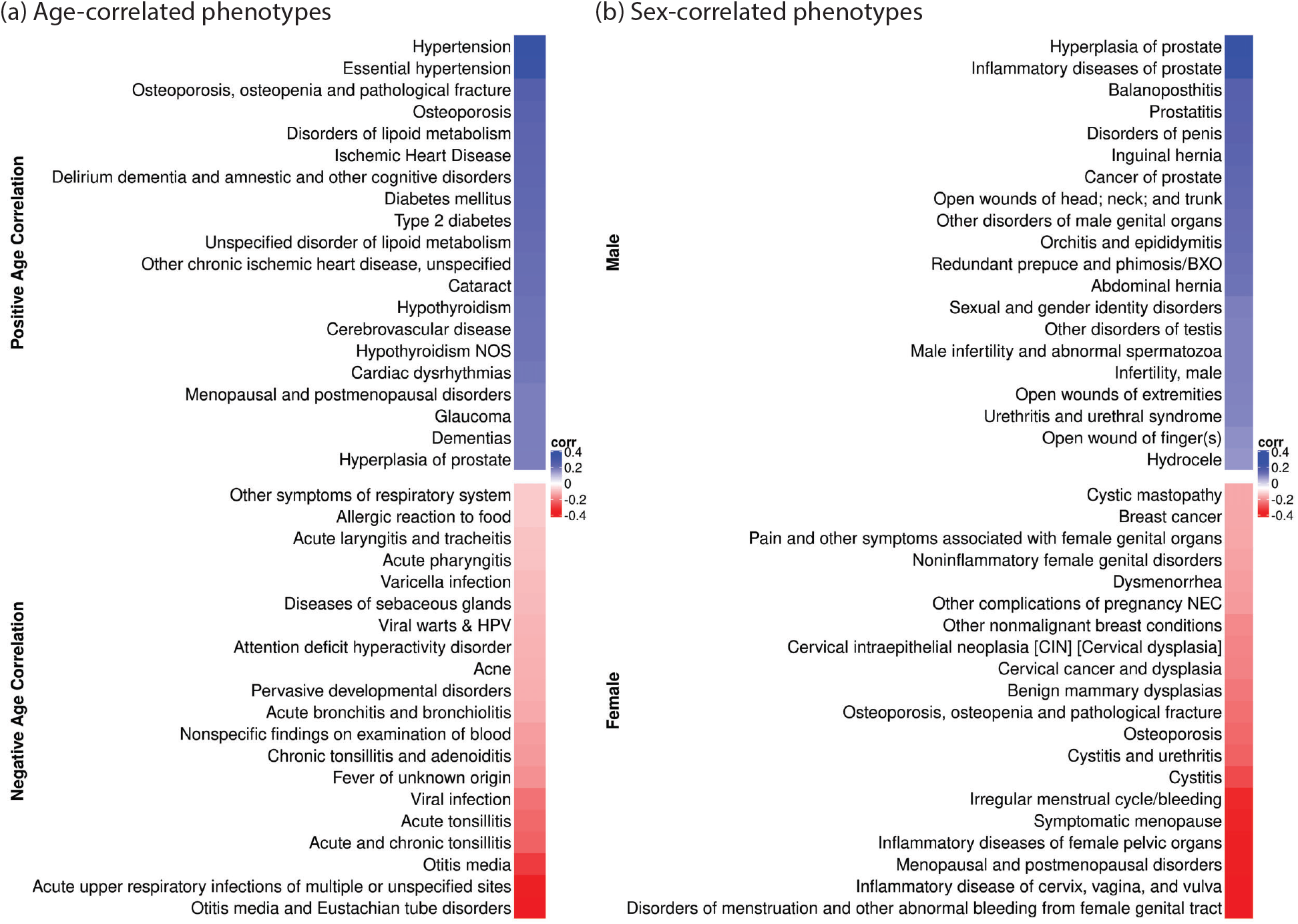
Top 20 phenotypes correlated with age and sex as predicted in our analysis. (**a**) The 20 phenotypes with the most positive (top) and most negative (bottom) correlations to patient age. (**b**) The top 20 sex-correlated phenotypes. For each panel, we computed Spearman correlations between MixEHR-G’s patient-topic mixtures 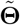 and patients’ (a) mean age or (b) sex, taking MALE as SEX=1.

**Figure 7:**
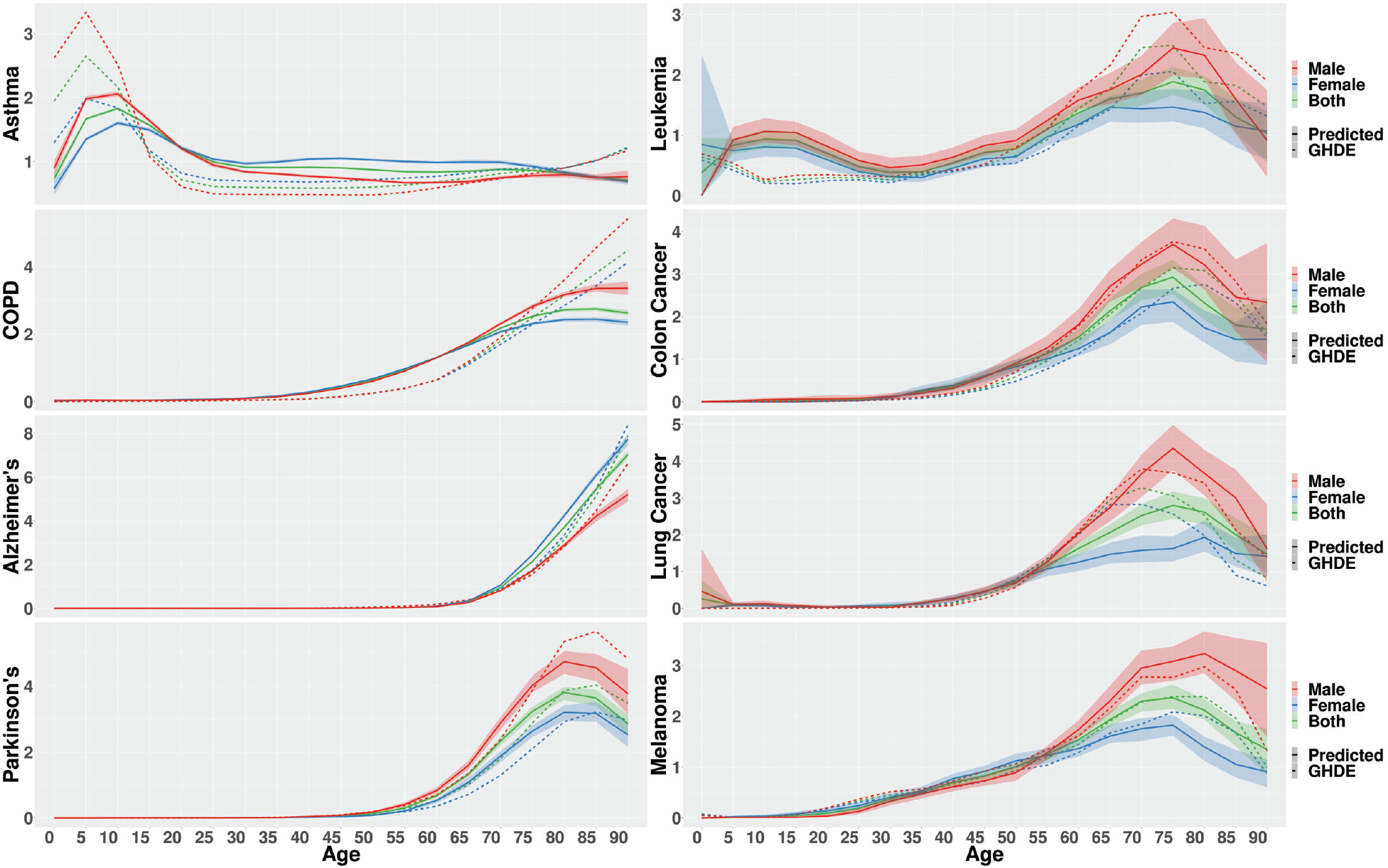
Estimates of relative phenotype prevalence. Based on the patient-phenotype mixtures 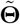, we computed phenotype prevalence stratified by age and sex for 8 diverse disease phenotypes with variable natural functions normalized by the mean estimate over time (**Methods**). We compared the predicted prevalences using MixEHR-G (solid lines) with the estimates from the Global Health Data Exchange (GHDE, dotted lines). 95% confidence intervals for MixEHR-G’s estimates were computed by bootstrapping with 100 replicates.

For a given age *t_age_*, we identified the set of patients 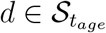 such that *T_min,d_* ≤ *t_age_* ≤ *T_max,d_*, where *T_min,d_* and *T_max,d_* denote the ages at which patient *d* respectively enters and exits the PopHR dataset. We then estimated the population phenotype prevalence at age *t_age_* as the mean predicted phenotype outcome over 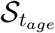:

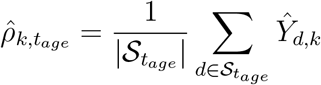

Recall that we only aimed to estimate relative, not absolute, prevalence over time. Given some predefined sequence of ages 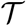 subscripted by *t*, we estimated the relative prevalence at time *t_age_* as

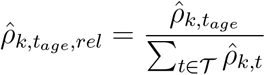

We compared our relative prevalence estimations to those from the IHME Global Health Data Exchange [47]. Again, we computed relative prevalence rates from the absolute prevalence rates supplied by the data exchange by normalizing *ρ_k_* over time.

## 5. Results

### 5.1. MixEHR-G’s phenotype topics are clinically meaningful

We qualitatively explored the inferred phenotype topics from both the PopHR and MIMIC-III datasets by examining the EHR features with the highest topic distribution probabilities (ϕ_k_’s) for 9 diverse chronic diseases covering a variety of medical specialities (**Fig.2**; **Supplementary Fig.** S4). We include top 5 features for all parent phenotypes and subphenotypes in the **Supplementary Materials**. We observe that in PopHR the phenotype topics appear to be predominantly defined by prescription (Rx) codes, which comprise 49.7%of feature tokens in the dataset (**Fig.2a**). Meanwhile, MIMIC-III phenotypes are dominated by ICD-9 codes (**Fig.2b**) despite that dataset consisting of 92.5%words from notes, 4.4%Rx codes, and only 1.6%ICD-9 codes. This highlights MixEHR-G’s ability to leverage potentially imbalanced EHR modalities to infer phenotype topics as opposed to relying solely on the ICD-9 signatures for each PheCode or drawing predominantly on the most prevalent features. In contrast, while sureLDA’s phenotype topics also diverge from the ICD-9 based priors, their top features tend to derive from each dataset’s most prevalent feature modality (prescription codes for PopHR and words from notes for MIMIC-III). Indeed, the top features sureLDA identifies for MIMIC-III are all words from notes, demonstrating sureLDA’s sensitivity to feature prevalence imbalances. (**Supplementary Fig.** S3).

Qualitatively, MixEHR-G consistently identifies clinically meaningful features for each phenotype topic (**Fig.2a**). For instance, risperidone and olanzapine are common antipsychotic medications used to treat schizophrenia (and to a lesser extent bipolar disorder), and furosemide is a common diuretic used for treatment of decompensated heart failure and cirrhosis, among other conditions. Likewise, MixEHR-G identifies meaningful associations between phenotype topics and lab results (**Supplementary Fig.** S2). For instance, the schizophrenia and bipolar topics exhibit high odds ratios for abnormal blood valproic acid, a common mood stabilizer, while the COPD topic is found to exhibit high odds ratios for abnormal urine chloride and potassium, reflecting the well known chronic compensatory metabolic alkalosis of these patients. Meanwhile, the COPD topic exhbits low odds ratios for abnormal troponin-1, a marker of ischemic cardiac injury, and NT-proBNP, a marker of heart failure - both diagnoses commonly ruled out in patients presenting with COPD exacerbation. Together these results suggest that while MixEHR-G is anchored to a prior defined by PheCodes, it is able to infer phenotype topics by leveraging other data modalities, including prescriptions and labs. Therefore, MixEHR-G’s phenotype topic distributions could be used to augment rule-based phenotyping algorithms and alleviate tedious manual phenotyping efforts.

We also observe qualitatively meaningful disease similarities and comorbidities based on the topic distributions (**Φ**) and patient-topic mixtures 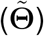 (**Fig.3)**. In particular, schizophrenia, bipolar disorder, and depression exhibit similar topic distributions with each other as well as with personality, anxiety, and mood disorders (**Fig.3a**). Phenotype pairs with known causative relationships also exhibit similar distributions, including diabetes and renal failure, heart failure and pulmonary edema, and cirrhosis and ascites. Interestingly, our analysis also revealed complex disease network connections among some phenotypes. For instance, cirrhosis is found to have a high degree of similarity with renal failure and nonhypertensive congestive heart failure, both of which also cause edema, as well as viral hepatitis, which also causes acute liver failure.

Relative similarity in terms of the patient-topic mixtures 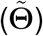 also replicates the well-known comorbidities among schizophrenia, mood disorders, and anxiety disorders, which have long been described in the medical literature [48–50] (**Fig.3b**). Other meaningful combinations include diabetes with ischemic heart disease and hypertension, as well as cirrhosis with ascites, biliary tract disease, and substance disorder. More interestingly, we observe comorbidities between bipolar disorder and hypothyroidism, concurrent with several recent studies positing a potential association between the two [51–53]. Similarly, the association between depression and myalgia/myositis supports the hypothesized clinical association between chronic pain/inflammation and depression [54, 55]. Notably, no two parent phenotypes have particularly strong associations (Spearman correlations < 0.4), reflecting the fact that even comorbid phenotypes like anxiety and mood disorders are very much independent entities.

### 5.2. Automated phenotyping using MixEHR-G’s topic mixture

A salient feature of MixEHR-G is that the K PheCode-guided topics are readily identifiable. Consequently, they can be directly used as *phenotype risk scores (PheRS)* [56]. Compared to alternative phenotyping algorithms, we found that MixEHR-G’s 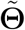 more closely aligns with validated rule-based phenotypes compiled by clinical experts at the Chronic Disease Surveillance Division of the Public Health Agency of Canada and elsewhere (**Methods**; **Fig.4**; **Table 1)** [57]. In particular, MixEHR-G’s scores exhibited significantly higher AUPRCs than the raw PheCodes or the recently published unsupervised phenotyping algorithms Multimodal Automated Phenotyping (MAP) [26] and sureLDA [30] for 9 out of 12 phenotypes evaluated (IHD, hypertension, HIV, diabetes, CPD, CHF, asthma, AMI, and ADHD) at level 0.05. For the remaining 3 phenotypes, either the PheCode already achieves near-perfect accuracy and thus leaves no room for improvement (i.e. schizophrenia, autism), or the phenotype’s prevalence is sufficiently low that MixEHR-G’s topic remains highly anchored to the PheCode prior (i.e. epilepsy).

**Table 1:** Accuracy metrics of MixEHR-G’s topic mixture scores 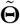 for identification of 12 diverse phenotypes. For computation of sensitivity, specificity, positive predictive value (PPV), negative predictive value (NPV), and F1 score, scores were thresholded to achieve NPVs of at least 0.95.

|  | Prevalence | AUROC | AUPRC | Sensitivity | Specificity | PPV | NPV | F1 |
| --- | --- | --- | --- | --- | --- | --- | --- | --- |
| ADHD | 0.02 | 1.00 | 0.94 | 1.00 | 1.00 | 0.88 | 1.00 | 0.93 |
| Acute MI | 0.03 | 0.84 | 0.48 | 0.70 | 0.98 | 0.52 | 0.99 | 0.60 |
| Asthma | 0.10 | 0.95 | 0.79 | 0.81 | 0.95 | 0.63 | 0.98 | 0.71 |
| Autism | 0.00 | 0.99 | 0.99 | 0.99 | 1.00 | 1.00 | 1.00 | 0.99 |
| CHF | 0.04 | 0.89 | 0.69 | 0.78 | 0.98 | 0.61 | 0.99 | 0.69 |
| COPD | 0.07 | 0.89 | 0.79 | 0.79 | 0.99 | 0.89 | 0.99 | 0.84 |
| Diabetes | 0.08 | 0.96 | 0.91 | 0.94 | 0.96 | 0.71 | 0.99 | 0.80 |
| Epilepsy | 0.01 | 0.60 | 0.16 | 0.20 | 1.00 | 0.67 | 0.99 | 0.31 |
| HIV | 0.00 | 0.92 | 0.76 | 0.84 | 1.00 | 0.57 | 1.00 | 0.68 |
| HTN | 0.19 | 0.94 | 0.88 | 0.86 | 0.95 | 0.81 | 0.97 | 0.83 |
| IHD | 0.09 | 0.94 | 0.81 | 0.91 | 0.95 | 0.64 | 0.99 | 0.75 |
| Schizo. | 0.01 | 0.99 | 0.99 | 0.99 | 1.00 | 1.00 | 1.00 | 0.99 |

MixEHR-G’s improvement over sureLDA was particularly notable despite the similarity between the two algorithms, suggesting that MixEHR-G’s multimodal topic design provides tangible benefits over sureLDA’s unimodal topics [30, 34]. The only phenotype for which sureLDA outperformed MixEHR-G was epilepsy, for which no phenotyping algorithm aligned particularly well with the rule-based labels (i.e. all AUPRCs were below 0.25). Nonetheless, MixEHR-G consistently equaled or outperformed the raw PheCode counts and MAP. Since both PheCodes and MAP only draw upon a handful of ICD-9 codes per phenotype, the superior performance of MixEHR-G further corroborates the benefit of leveraging the vast amount of additional information in patients’ EHRs.

To investigate the concordance and discrepancy between MixEHR-G’s inferred topics and the existing phenotyping rules, we focused within the PopHR dataset on chronic obstructive pulmonary disease (COPD), a notoriously difficult phenotype to define with rules (**Fig.5**; **Supplementary Fig.** S5) [58]. We first examined the top 25 codes under the COPD topic’s distribution ϕ_COPD_ (**Fig.5a**; **Supplementary Fig.** S5). Four out of eight ICD codes that constitute the COPD rule were present among the top 10 features under the COPD topic distribution. However, MixEHR-G also incorporates other meaningful features including prednisone (commonly used for treatment of COPD exacerbations), salbutamol (used for asthma and COPD maintenance but more commonly COPD), salmeterol (also used for asthma and COPD maintenance but more commonly COPD), and tiotropium (used almost exclusively for COPD maintenance). Indeed, MixEHR-G identifies COPD patients using all available information in the EHR as opposed to just the ICD-9 codes defined by the rule.

We further examined the top 10 rule-negative patients with the highest MixEHR-G COPD PheRS scores (**Fig.5b** left panel) as well as the top 10 rule-positive patients with the lowest MixEHR-G COPD PheRS scores (**Fig.5b** right panel). Many of the patients in the former category have numerous observations of prednisone, fluticasone, salbutamol, salmeterol, theophylline, or tiotropium, suggesting that they may indeed be COPD patients (i.e. potential false negatives for the rule). Conversely, many low-scoring patients in the latter category exhibit no evidence of any relevant prescriptions, making them dubious COPD cases (i.e. potential false positives). These results suggest that the predefined disease validation rules may be too rigid since no ICD code or prescription is adequately sensitive or specific. On the other hand, MixEHR-G provides richer and largely complementary information to help re-define these rules in a data-driven manner. To formally assess this, we would need chart-reviewed gold-standard labels, which are not available from our current data.

### 5.3. Estimating disease prevalence rates in Quebec using MixEHR-G’s topic mixture

We used MixEHR-G’s patient-phenotype mixtures 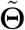 to identify which phenotypes are most strongly associated with age and sex (**Fig.6)**. We observe that hypertension and osteoporosis are strongly positively associated with age, whereas otitis media and tonsilitis are strongly negatively associated. Likewise, pathologies of the prostate are associated with male sex, whereas breast and cervical pathologies are associated with female sex. These simple proof-of-concepts paved the way for further exploratory epidemiological analyses.

We used 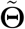 to investigate the relative prevalence for 8 diverse phenotypes stratified by age and sex, which we compare to gold-standard estimates from the Global Health Data Exchange (GHDx) [47] (**Fig.7**; **Methods**). We observe high concordance between our estimates and the gold-standard. Consistent with the gold-standard, our analysis indicates that asthma peaks in prevalence around age 10; leukemia and lung cancer around ages 65-75; Parkinson’s disease, colon cancer, and melanoma around ages 75-85, and COPD and Alzheimer’s disease just increase monotonically with age. Likewise, MixEHR-G’s estimates support the inference that asthma, COPD, Parkinson’s disease, colon cancer, and melanoma are significantly more prevalent in men, whereas Alzheimer’s disease (AD) predominates in women, likely reflecting increased survivorship among female Alzheimer’s patients. These diseases were arbitrarily chosen as a proof-of-concept based on our knowledge about the diverse onsets of these diseases and the availability of prevalence data from GHDE. We leave a more comprehensive analysis of the entire phenome as future work.

## 6. Discussion

The effective application of machine learning algorithms to EHR data will lay the foundation for the development of modern digital medicine. One important and fruitful application is predicting a patient’s disease phenotypes using a supervised learning model such as logistic regression or neural network [59]. However, supervised learning methods are not scalable for simultaneously predicting thousands of phenotype labels [18]. Modelbased unsupervised methods such as non-negative matrix factorization [60], autoencoder [31], and topic modeling [34] are at the other end of the spectrum. While the latent factors or topics inferred by these unsupervised methods can provide clinical insights, they are not identifiable as inferred topics cannot be directly mapped to known phenotypes. Moreover, while supervised topic models such as MixEHR-S [61] can simultaneously infer topic distributions and fit a predictive function of a target disease, they are not scalable to predicting multiple disease labels simultaneously. Therefore, an efficient method is needed to achieve simultaneous phenotype inference while providing interpretable topic distributions over heterogeneous EHR data.

In this study, we present a PheCode-guided topic modeling algorithm called MixEHR-G to simultaneously model up to 1515 well-defined phenotypes as a function of highdimensional, multimodal EHR data. MixEHR-G is highly scalable due to its use of memory-efficient sparse matrices and time-efficient variational Bayesian inference implemented with parallelization in C++. Indeed, it can perform inference on 1.3 million patient records containing a total of 70.7 million non-zero feature observations over 24,192 unique features in ~10 hours on a 2.2 GHz processor with 10 cores, using a maximum of 60 MB of RAM and 130% CPU (1.3 cores).

In proof-of-concept analyses, we observe that the MixEHR-G-inferred topics are well-aligned with the phenotypes they represent and complementary to rule-based phenotyping algorithms. We used MixEHR-G trained on the PopHR database to derive meaningful insights about disease similarities, comorbidities, and prevalences in the Quebec population [62].

We envision two major applications for the trained MixEHR-G model. The first application is phenotype labeling for cohort selection or clinical research. MixEHR-G more closely aligns with expert-curated phenotype rules than other recently published phenotyping methods including MAP and sureLDA across 9 diverse phenotypes (**Fig.4)**. Our case study on COPD suggests that effectively incorporating prescription data produces more informative rules (**Fig.5)**. For a large-scale downstream study that requires labels across many phenotypes, such as a Phenome-Wide Association Study (PheWAS), MixEHR-G’s patient-topic mixtures 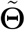 can be directly used as continuous phenotype risk scores (PheRS) [56], obviating the need for labor-intensive chart review or rule generation.

For a study focusing on a single phenotype, MixEHR-G’s topic distributions ϕ can be further refined with subdivided groups of specific EHR features for sub-phenotyping [56]. When associating these sub-phenotypes with genotype data using population data such as UK Biobank [63], we can investigate the genetic epidemiology of the refined subphenotypes and gain insights into the diverse outcomes of the same primary disease (e.g., drug response, survival time). These phenotype labels can then be used for downstream epidemiological research or to identify cohorts for inclusion in a clinical trial.

For patient outcome classification, the inferred phenotype topic probabilities 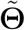 can be treated as phenotype risk scores or PheRS [56] derived from the training patient dataset. As a post-processing step, they can then be incorporated as strongly predictive features into a supervised classifier for applications such as short-term mortality prediction [34, 64], sequential diagnosis prediction [34], and first-time drug usage (e.g., insulin regime adoption by diabetic patients [61]). Given the interpretability of the topics, we can then use them to understand the target outcome based on their corresponding importance scores derived from the trained classifiers.

MixEHR-G’s second major application is to derive population-level insights about phenotypes. In this study, we validated the use of MixEHR-G’s patient-topic mixtures 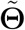 to infer relative phenotype prevalence stratified by age and sex. For prevalence estimation specifically, MixEHR-G’s estimates can either elucidate the general shape of a prevalence function over a stratifying variable such as age, or be calibrated using a small set of gold-standard labels and a validated risk estimation method such as SCORNET [65] to produce valid absolute prevalence estimates. Moreover, MixEHR-G’s outputs can identify potential epidemiological directions for more rigorous follow-up. For instance, our analysis of associations in ϕ supports the recently hypothesized comorbidity between bipolar disease and hypothyroidism (**Fig.3b**); ascertaining the nature of this association could be an interesting subject for further investigation. On the other hand, we can incorporate the prior knowledge of the phenotypic prevalence by sex and age (e.g., from Global Health Data Exchange) into the Bayesian topic prior for each patient (i.e., the Dirichlet topic hyperparameters). Therefore, depending on the age and sex of patient *d*, we will have a different set of prior weights over his/her topic mixture membership θ_*d*_. This will presumably improve phenotyping accuracy especially for small sample size and sparse observation in the EHR data.

As a future direction, we will extend MixEHR-G to model 1) continuous data modalities, and 2) time. In its current form, MixEHR-G must threshold continuous variables such as lab tests into categorical values, and it rolls up observations over time into feature counts to infer longitudinal patient parameters. While many continuous clinical variables can be safely reduced to low-dimensional categoricals without significant loss of information (i.e. certain lab results being represented as ‘normal’, ‘high’, or ‘low’), many other variables (i.e. age) cannot be easily reduced in this way. Moreover, being able to model the progression of patients’ phenotype topics over time could yield valuable insights about longitudinal risk, the natural course of diseases, and causal relationships. We leave these as future extensions for the MixEHR model family.

We also envision MixEHR-G being modified to handle multi-center data such that it can simultaneously incorporate EHR data across medical centers. In MixEHR-G’s current form, incorporating multi-center data would be challenging with potential biases given meaningful differences in not just patient populations but also how diseases are coded and treated. Being able to leverage commonalities across centers in a manner similar to mixed effects models could yield powerful insights on a scale larger than any individual medical center.

## 7. Conclusion

In summary, by modeling multimodal phenotypes with the guided topic modeling strategy, MixEHR-G achieves accurate simultaneous phenotype predictions for up to 1515 phenotypes and improved interpretability of phenotype topics. Altogether, we believe that MixEHR-G will enable more efficient use of EHR data toward the goals of personalized digital medicine and precision public health.

## Ethics

The use of the PopHR data for this research was approved by the McGill Faculty of Medical and Health Sciences Institutional Review Board.

## Acknowledgements

Y.L. is supported by Natural Sciences and Engineering Research Council (NSERC) Discovery Grant (RGPIN-2019-0621), Fonds de recherche Nature et technologies (FRQNT) New Career (NC-268592), and Canada First Research Excellence Fund Healthy Brains for Healthy Life (HBHL) initiative New Investigator start-up award (G249591). D.B. is supported by a Canada Research Chair (Tier 1) in Health Informatics and Data Science. The authors declare no conflicts of interest.

## Author contributions

Y.A., D.B., and Y.L. conceived the study. Y.A. and Y.L. developed the methodology and implemented the software. Y.A. ran the majority of the data analyses with help from Y.Z. and A.V.. Y.A., D.B., and Y.L. analyzed the results. Y.A. and Y.L. wrote the initial draft of the manuscript with critical comments from D.B. All of the authors reviewed and wrote the final version of the manuscript.

## Supplementary Materials

### S1. Supplementary Figures

**Figure S1:**
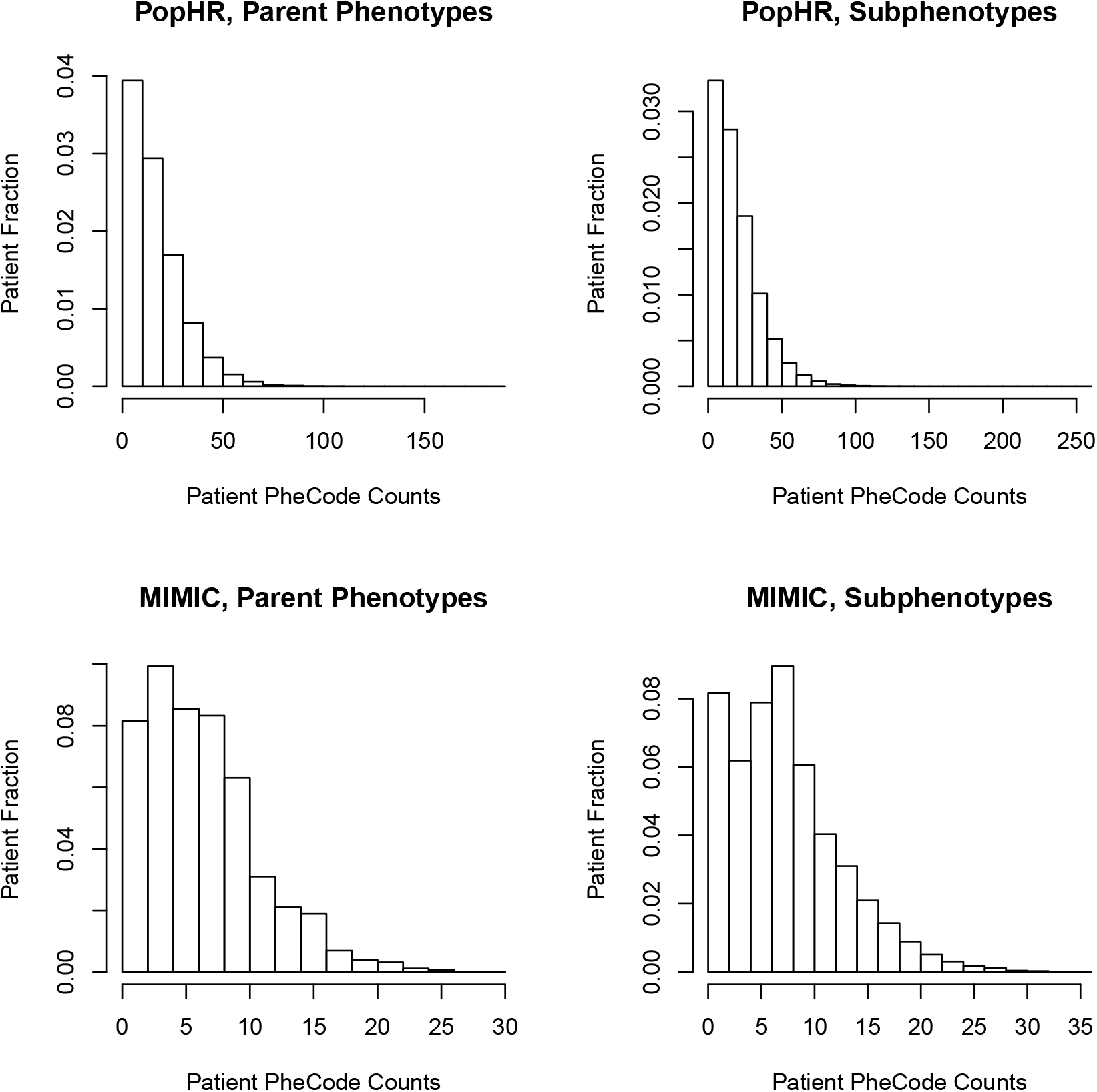
Distribution of PheCode anchors per patient (including both 3-integer and 3-integer+1-decimal codes) for the PopHR and MIMIC-III datasets.

**Figure S2:**
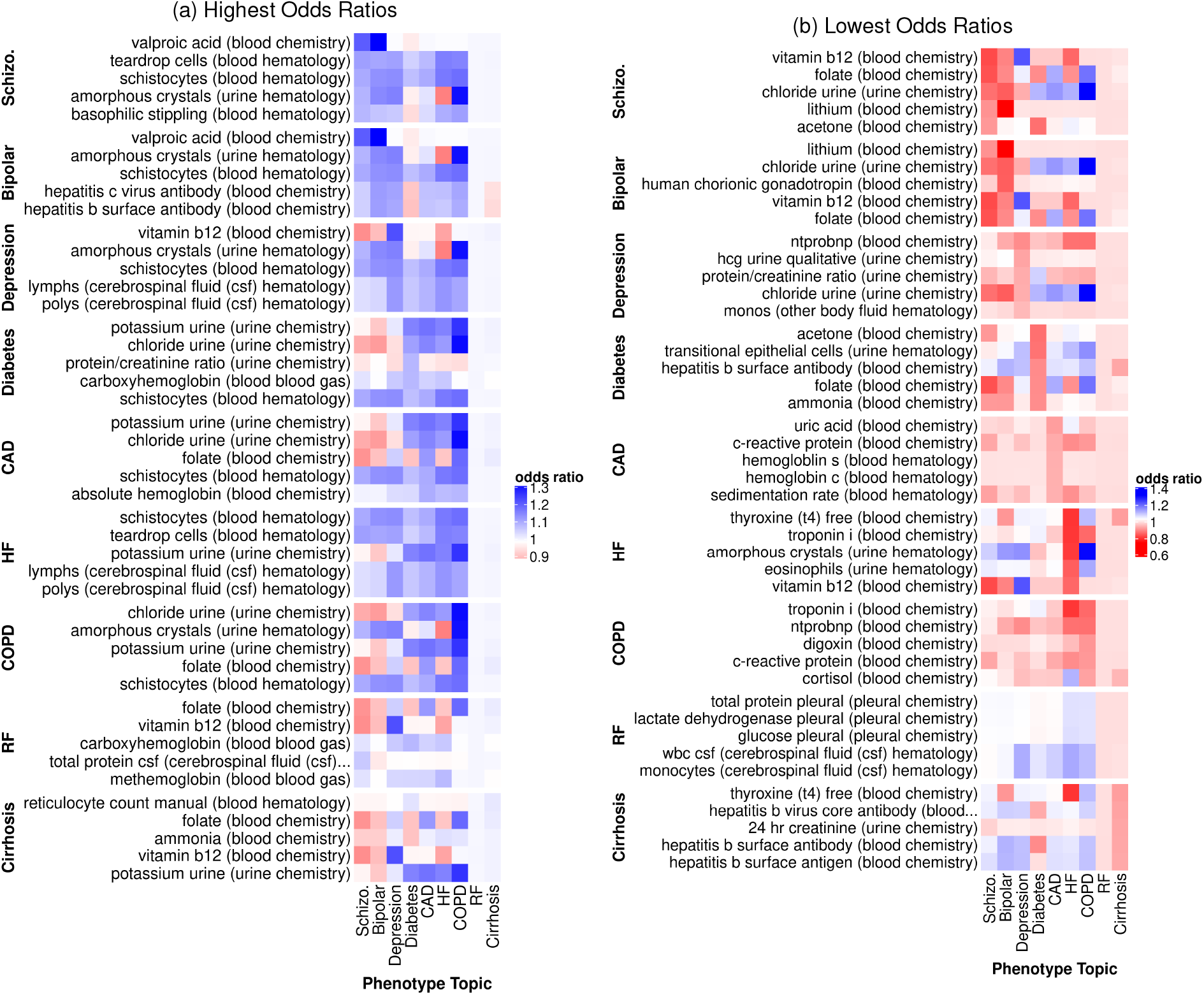
Lab tests with the (a) highest and (b) lowest odds ratio of abnormal result for each of 9 diverse disease phenotypes as ascertained by MixEHR-G trained on the MIMIC-III dataset.

**Figure S3:**
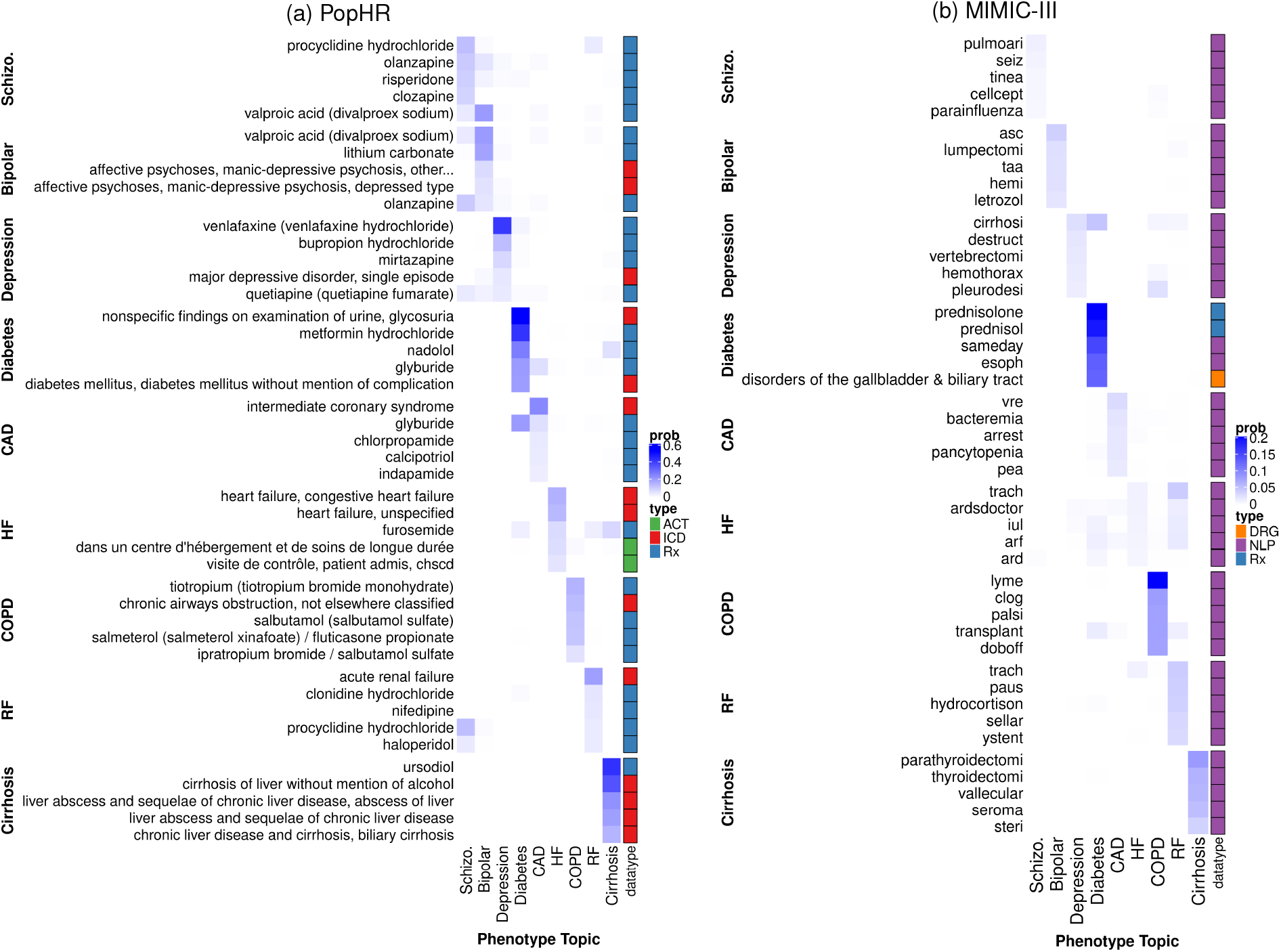
Top 5 features for each of 9 diverse disease phenotypes as ascertained by sureLDA, for comparison with **Fig.** 2. We present results for two datasets: (a) PopHR, which has 3 data modalities, and (b) MIMIC-III, which has 5.

**Figure S4:**
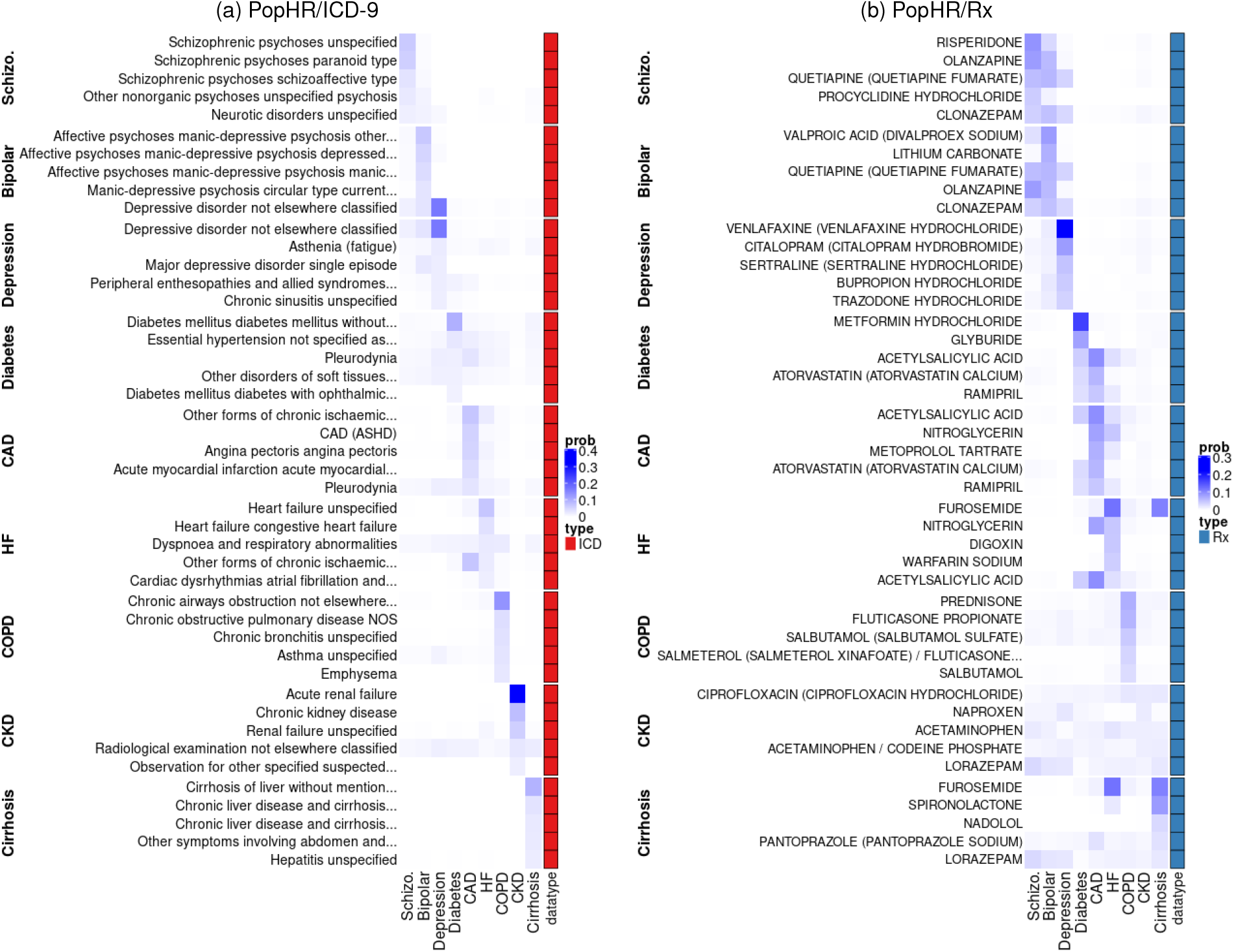
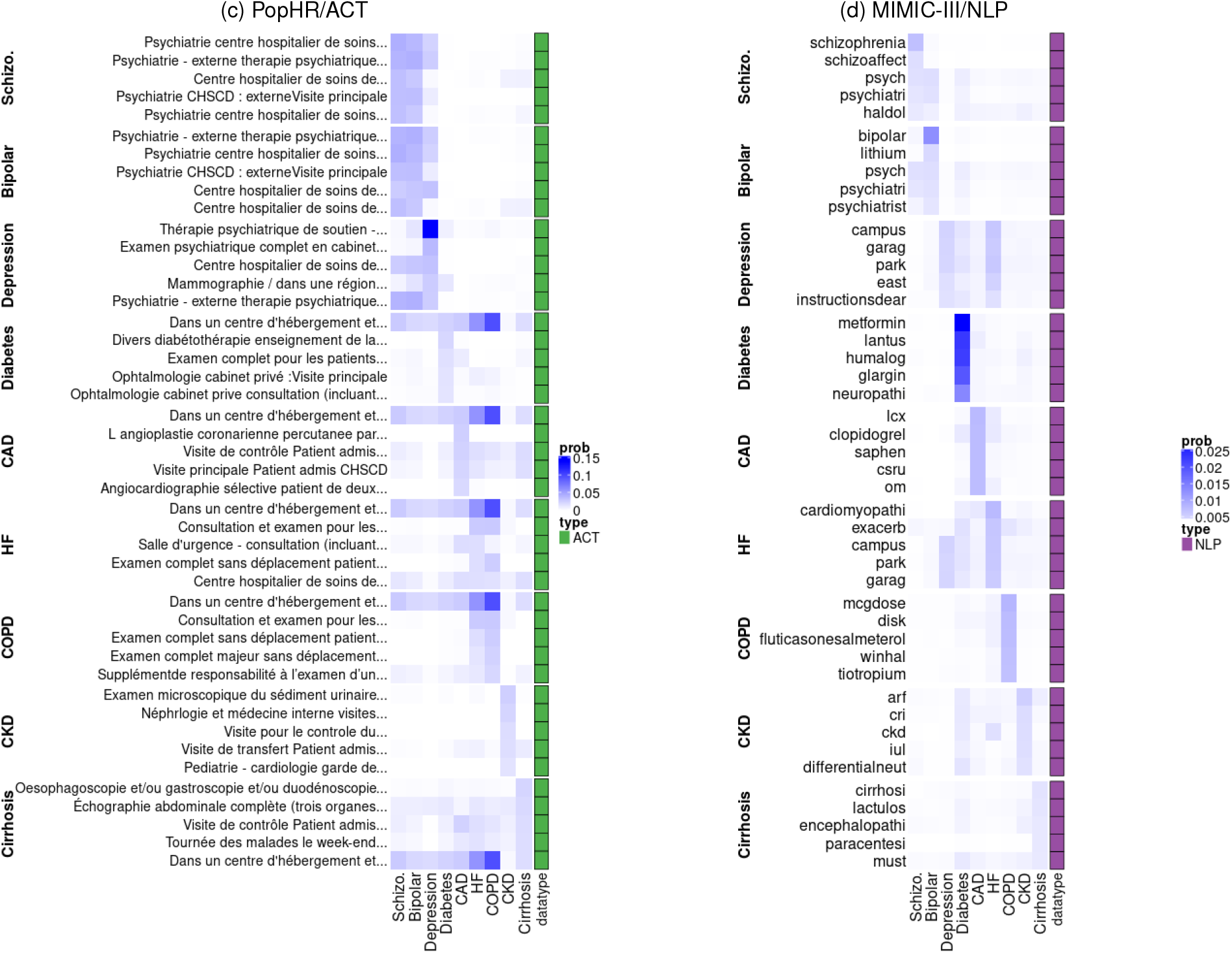
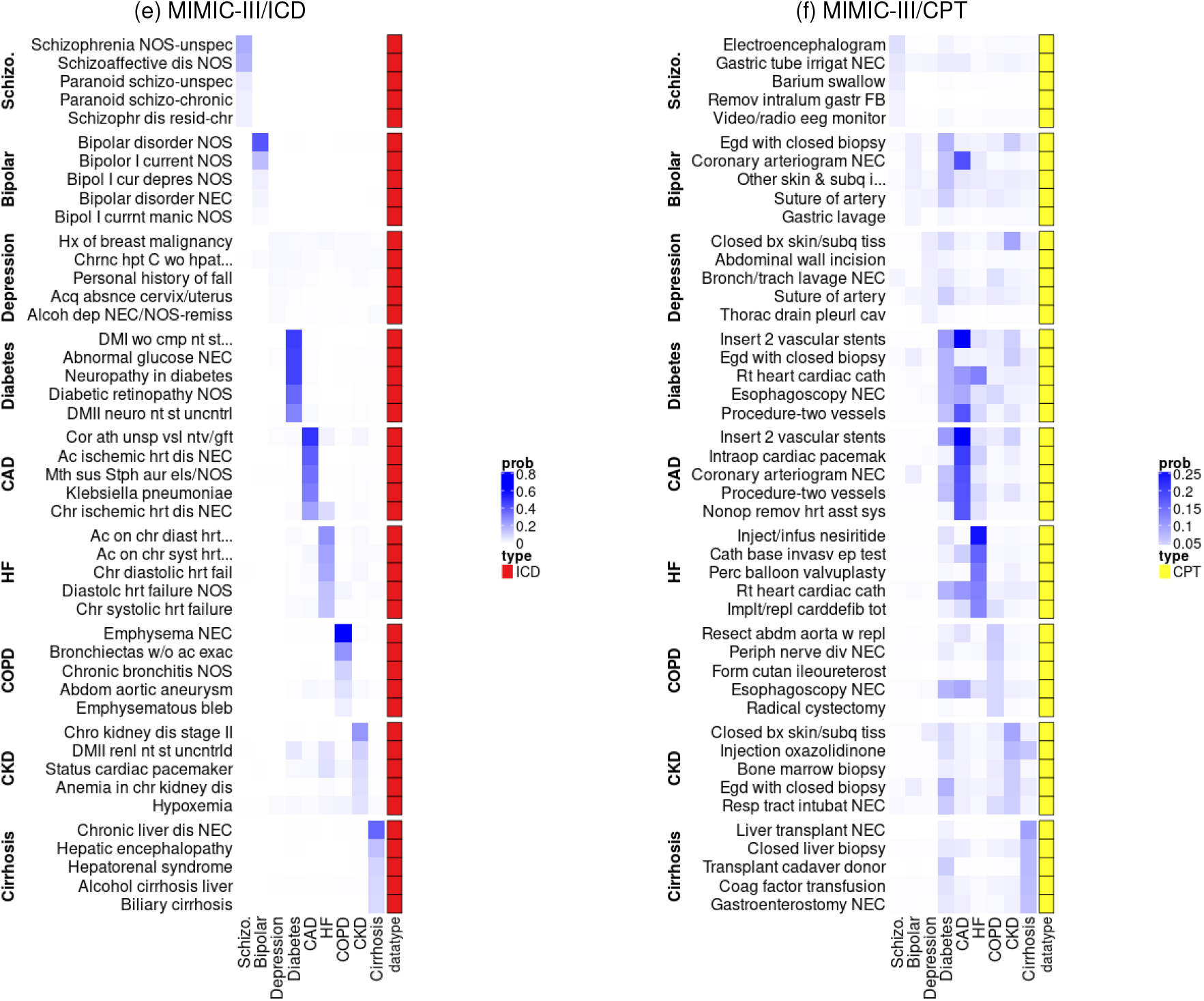
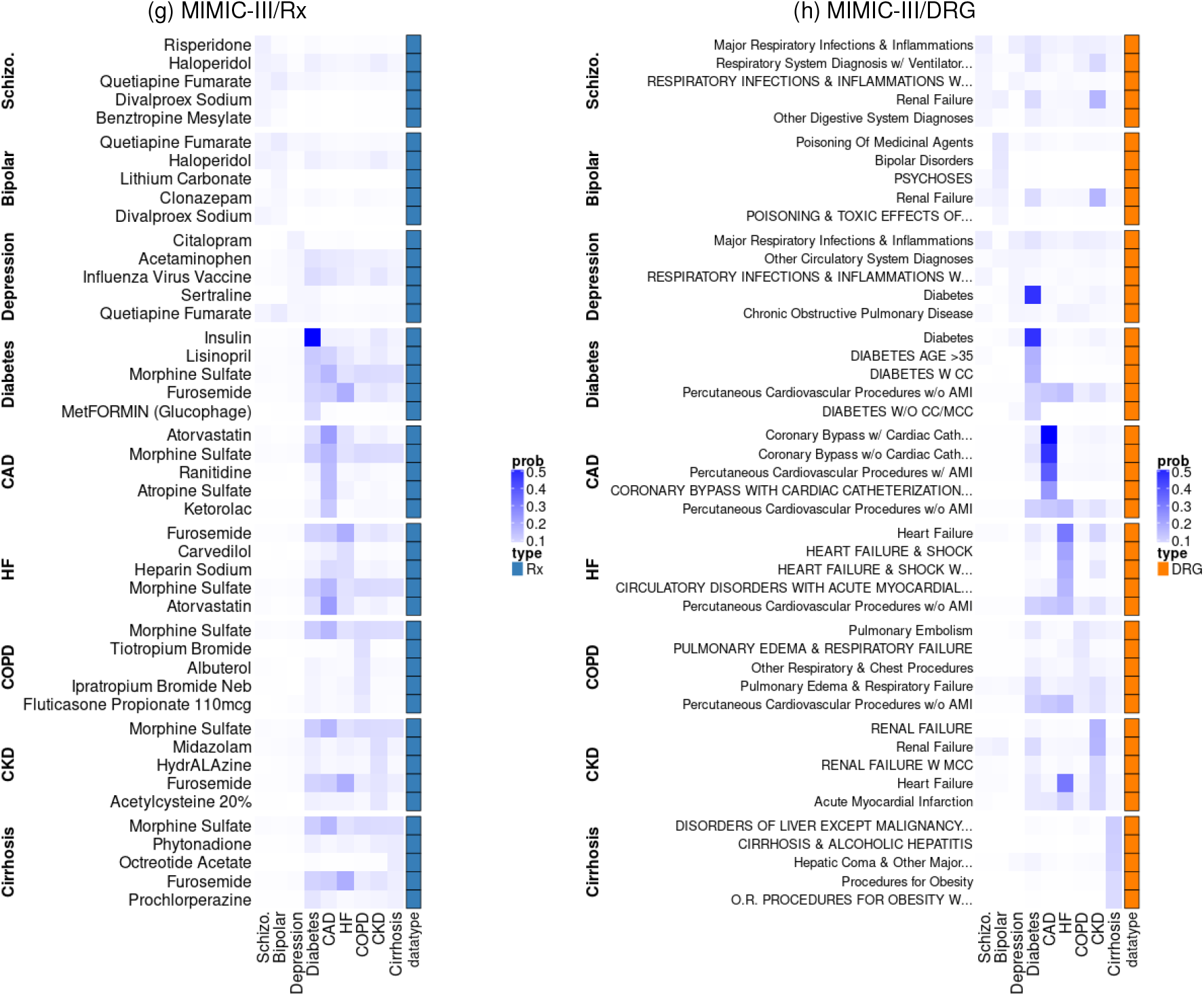
Top 5 features stratified by data modality for each of 9 diverse disease phenotypes as ascertained by MixEHR-G. We present results for two datasets: (a-c) PopHR, which has 3 multinomial data modalities (ICD-9, ACT, and Rx), and (d-h) MIMIC-III, which has 5 (NLP, ICD-9, CPT, Rx, and DRG). Note that while MIMIC-III also contains lab observations and results, MixEHR-Gdoes not model these modalities as multinomials, and thus their corresponding distributions cannot be interrogated in this way.

**Figure S5:**
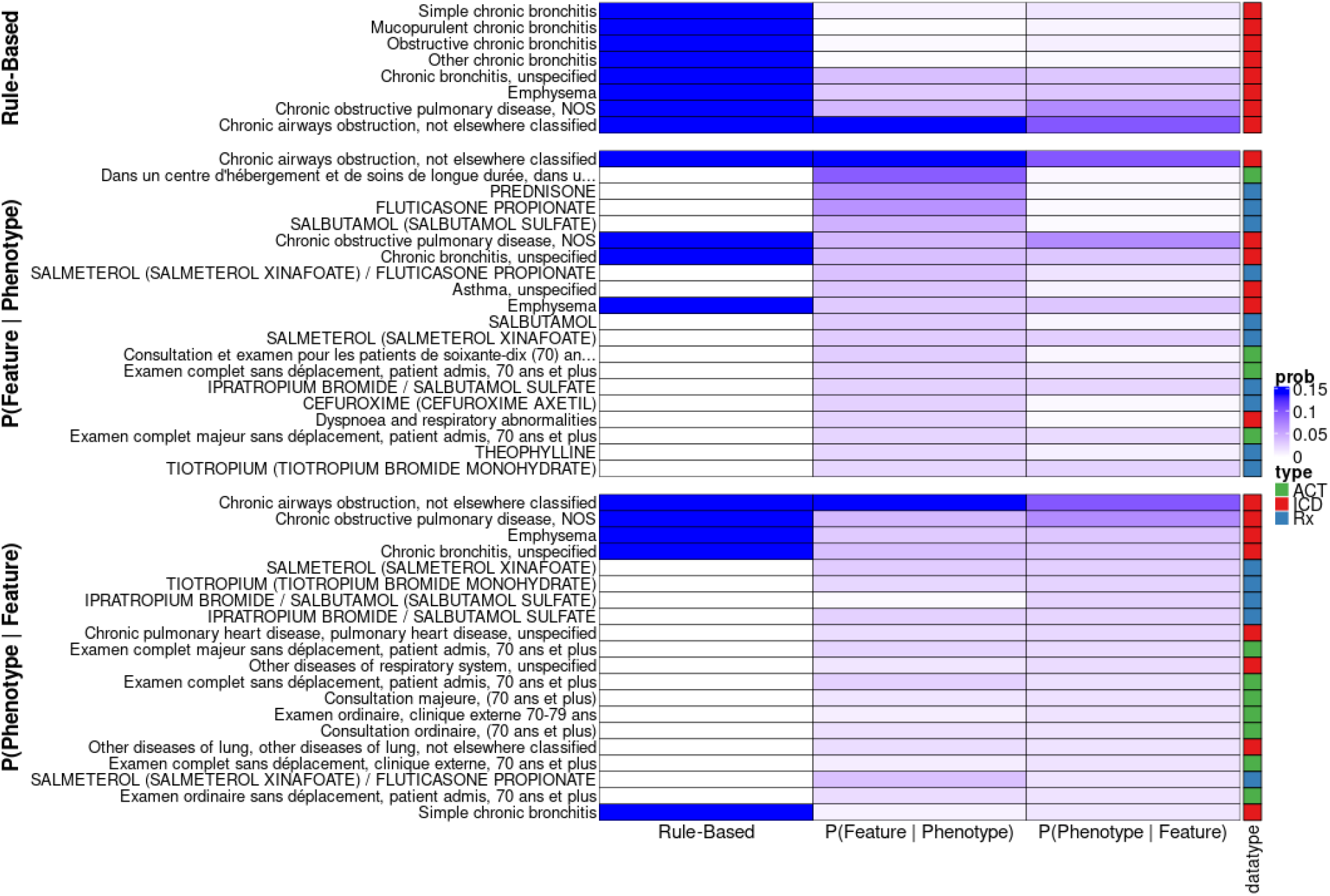
Top 25 features with the highest (1) sensitivities {P(Feature | Phenotype)} and (2) positive predictive values {P(Phenotype | Feature)} for COPD based on MixEHR-G’s trained topic distributions ϕ, as compared to features included in the rule-based COPD algorithm.

